# Ubr1-induced selective endo-phagy/autophagy protects against the endosomal and Ca^2+^-induced proteostasis disease stress

**DOI:** 10.1101/2021.10.05.463117

**Authors:** Ben B. Wang, Haijin Xu, Sandra Isenmann, Cheng Huang, Xabier Elorza-Vidal, Grigori Y Rychkov, Raúl Estévez, Ralf B. Schittenhelm, Gergely L. Lukacs, Pirjo M. Apaja

## Abstract

The defence mechanisms against endo-lysosomal homeostasis stress remain incompletely understood. Here, we identify Ubr1 as a protein quality control (QC) ubiquitin ligase that counteracts proteostasis stress by enhancing cargo selective autophagy for lysosomal degradation. Astrocyte regulatory cluster membrane protein MLC1 mutations increased intracellular Ca^2+^ and caused endosomal compartment stress by fusion and enlargement. Endosomal protein QC pathway using ubiquitin QC ligases CHIP and Ubr1 with ESCRT-machinery was able to target only a fraction of MLC1-mutants for lysosomal degradation. As a consequence of the endosomal stress, we found an alternative QC route dependent on Ubr1, SQSTM1/p62 and arginylation to bypass MLC1-mutants to endosomal autophagy (endo-phagy). Significantly, this unfolded a general biological endo-lysosomal QC pathway for arginylated Ubr1-SQSTM1/p62 autophagy targets during Ca^2+^-assault. Conversely, the loss of Ubr1 with the absence of arginylation elicited endosomal compartment stress. These findings underscore the critical housekeeping role of Ubr1-dependent endo-phagy/autophagy in constitutive and provoked endo-lysosomal proteostasis stress, and link Ubr1 to Ca^2+^-homeostasis and proteins implicated in various diseases including cancers and brain disorders.

## Introduction

Disease-causing mutations, as well as perturbations in intracellular Ca^2+^ homeostasis, may result in protein misfolding and/or accumulation in the cytosol or membrane organelles, mitochondrial oxidative damage^1^, the endoplasmic reticulum (ER) stress or lysosomal swelling and exocytosis^2^ that have been linked to numerous Mendelian disorders, neurodegenerative diseases, and cancers. To protect protein homeostasis (proteostasis), cells have developed spatial or organelle-specific protein QC mechanisms^3^. At the ER, QC mechanisms recognize various degradation signals (degrons) in luminal, transmembrane or cytosolic segments of misfolded membrane proteins as part of the ER-associated degradation (ERAD) and ER-phagy^4, 5^. The cytosolic PQC preferentially relies on the coordinated actions of molecular chaperones, ubiquitin (Ub) conjugation machinery, proteasomes (Ub-proteasome systems) and chaperone-mediated autophagy^3^.

Conformationally defective cell surface membrane proteins generated either in situ or escaped from the ER QC are recognized by the peripheral PQC machinery. Primarily, increased post-translational conjugation of ubiquitin (ubiquitination) in non-native plasma membrane (PM) proteins is a signal for ESCRT (endosomal sorting complex required for transport)-dependent targeting to the multivesicular body (MVB) and lysosomal degradation^6–10^. The role of peripheral PQC is to maintain native protein composition and intra- and inter-organelle homeostasis during membrane dynamics between Golgi, the plasma membrane (PM)- and endo-lysosomal compartments. The peripheral PQC is less studied, and only three QC E3 Ub-ligases have been implicated in the clearance of non-native PM and endosomal membrane proteins to lysosomal proteolysis. The C-terminus of Hsc70-Interacting Protein (CHIP), Nedd-4 (and its yeast homologues Rsp5), as well as the RFFL, represent both chaperone-dependent and -independent PQC mechanisms^6, 7, 9, 11, 12^.

Additional post-translational modifications by sumoylation, phosphorylation or arginylation with ubiquitination may enrich PQC signalling in a substrate-specific manner^13–16^. From these, arginylation has a role outside QC as N-terminal degron for n-recognin type proteins in fast proteasomal degradation^17^. Previous research also found that N-terminal arginylation of BiP (HSPA5, GRP78) released from the ER and cytosolic misfolded soluble proteins can be cleared by autophagy upon proteasomal inhibition or cytokine-induced oxidative stress^18–21^. In the current study, we found new roles for n-recognins in PQC during Ca^2+^-assault, endosomes and cargo selective autophagy, different from their previous roles at the N-degron pathway.

The formation of phagophores, the newly formed membranes that engulf autophagy cargo, has been demonstrated at Rab11+ early endosomes^22^. Instead, endosomal autophagy cargo recognition and clearance mechanisms for misfolded endosomal membrane proteins or in endosomal stress QC remain undefined. To address these questions, we start the investigation by examining the endosomal PQC activity using disease-associated regulatory membrane protein MLC1 (megalencephalic leukoencephalopathy with subcortical cyst) and end the study addressing separately endosomal compartment PQC during Ca^2+^-stress.

MLC1 is a unique part of the astrocyte PM macromolecular regulatory signalling cluster, pertinent for astrocytes homeostasis, motility/morphology^23–25^, inflammatory responses^26^, signalling and ion homeostasis regulation of the brain extracellular space^27–29^. The cluster encompasses integral membrane proteins including Na^+^/K^+^-ATPase, TRPV4 cation and ClC-2 chloride channels, EGF receptors and GlialCAM adhesion molecules^30–33^. GlialCAM is required for the correct MLC1 cell surface entry and tetherin^34^. At the cell surface, MLC1/GlialCAM regulates the diffusional partitioning and endosomal dynamics of the cluster^34^, as well as the activity of ClC-2^32^, gap junctions^25, 28^ and Na^+^-K^+^-ATPase^35, 36^. Disease mutations in MLC1 or GlialCAM and/or altered expression or compromised PM tethering of the cluster result in MLC1-dependent loss of endo-lysosomal organellar identity and impaired cargo sorting^34^. Thus, MLC1 has far-reaching consequences on astrocytic and neuronal proteostasis with the incompletely understood molecular mechanism.

Here we show that Ubr1 is a key player in the biological stress PQC to protect the endosomal compartment proteostasis. We found that disease-causing MLC1 variants cause endo-lysosomal and intracellular Ca^2+^-stress are partly offset by E3 Ub-ligases CHIP and Ubr1 to facilitate mutants ESCRT-mediated lysosomal proteolysis. Remarkably, as an alternative triage mechanism, Ubr1- and SQSTM1/p62-dependent selective endosomal autophagy (endo-phagy) cleared ubiquitinated and arginylated MLC1 clients. Further evidence showed Ubr1/SQSTM1/p62-axis to be a wider QC mechanism during Ca^2+^-load involved in cargo selective dual ubiquitin/arginine-client autophagy to maintain proteostasis. Our study connects Ubr1 and ubiquitin/arginine targets to a previously unidentified form of PQC and potentially to many human proteostasis diseases.

## Results

### Mutant MLC1s are targeted for ubiquitination

GlialCAM functions as a chaperone-like QC factor to MLC1 in the ER ensuring its cell surface expression^34^. Disease-causing mutations in MLC1 or GlialCAM have reduced cellular and PM turnover, and the same enlarged endosomal compartment phenotype^34^. The expression of regulatory MLC1 appears to be critical in the PM cluster. Therefore, we set out to examine characterized^34^ mutations possessing gradual decline (P92S:mild, S280L:moderate, C326R:severe) in their PM turnovers and expression (^34^, Fig. 1A^37^) using doxycycline-inducible HeLa and astrocytic U251N cells, immunoprecipitations (IP) and live-cell surface (cs)-ELISA for the PM-endosomal kinetics (^31, 32, 34^, Methods). The expressed MLC1-wt was comparable to that of the endogenous MLC1 in U251N cells (Fig. S1A). Proteasomal inhibition (MG132, 2h) unmasked increased total poly-Ub of the IPed P92S relative to the wt (Fig. S1B), suggesting that misfolding increased Ub and reduced steady-state expression. To examine specifically the post-Golgi turnover, we expressed MLC1-P92S in ts20 cells harbouring the thermolabile E1 Ub-activating enzyme. Inactivation of the E1 enzyme at 40°C (Fig. S1C) not only eliminated the MLC1-P92S ubiquitination (Fig. S1D) but also decreased four-fold its PM turnover from a half-life (T_1/2_) of ∼1.5h to ∼4h, approaching that of the wt (Fig.1B). This suggested that ubiquitination of the ER QC escaped MLC1-P92S accelerated the PM turnover. As support, P92S endosomal internalization was inhibited by >50% in ts20 cells than in control E36-wt cells (Fig. S1E). The MLC1-C326R had pronounced PM destabilization (T_1/2_ ∼0.5h) and steady-state expression defect (Fig.1B)^31^. These results indicate that the significantly reduced steady-state expression of mutants is attributed to their accelerated turnover in post-Golgi compartments.

**Figure 1.**
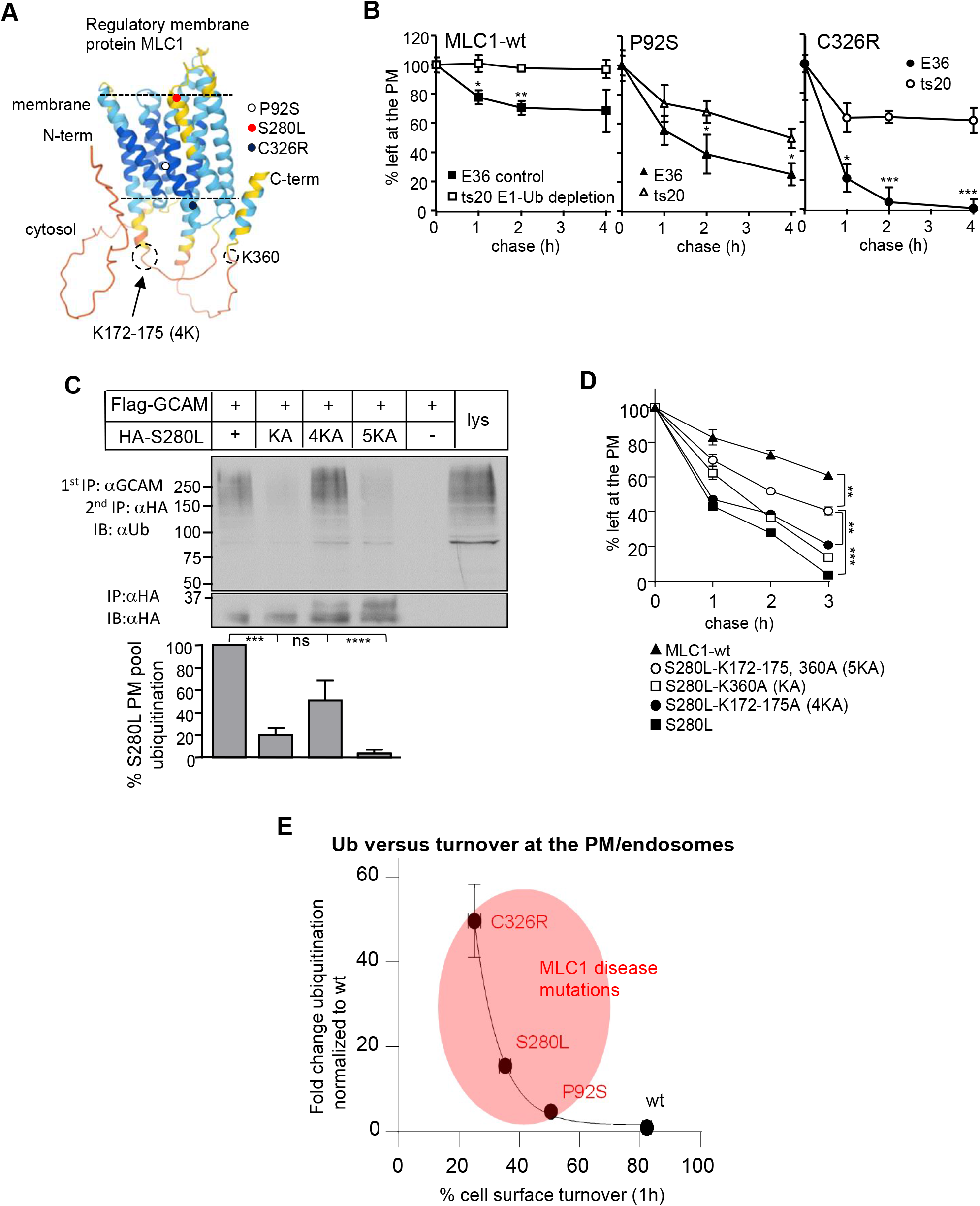
Astrocyte membrane regulatory cluster MLC1 disease variants are ubiquitinated at the PM-endosomal compartment. **A)** Predicted regulator membrane protein MLC1 Alphafold^37^ structure with indicated disease mutations (P92S, S280L and C326R) used in this work and ubiquitin lysine (K) acceptor sites used in Fig. 1C-D. **B)** Lack of the temperature-sensitive (ts) E1 ubiquitin (Ub)-activating enzyme decreased the PM turnover of MLC1 variants. MLC1-wt and disease associated mutations P92S and C326R were transiently expressed in E36 control and ts20 cells and the E1 was inactivated at 40°C for 3h to remove Ub-conjugation. Live-cell surface (cs)-ELISA for HA-MLC1 and chase for indicated times were used to measure turnover as described in Methods. **C)** Ub-acceptor lysine mutations decrease the PM-endosomal ubiquitination of misfolded MLC1-S280L. The MLC1/GlialCAM complex was first cell surface immunoprecipitated (cs-IP) using GlialCAM Ab and the second IP was performed with HA Ab for MLC1 after denaturation. Flag-GlialCAM was transiently expressed in stable inducible HA-MLC1 HeLa^31, 32^. Ubiquitin acceptor K was mutated (KA=K360, 4KA=K172-175, and 5KA=KA+4KA, see also Fig.1A). The ubiquitination was normalized to the S280L amount (100%) at the PM. GCAM, GlialCAM; lys, lysate; α, anti. **D)** Ub-acceptor lysine mutations decrease the PM turnover of MLC1-S280L measured using cs-ELISA as in A. **E)** Correlation of MLC1 disease variants (P92S, S280L, C326R) PM turnover^34^ to the relative ubiquitination at the PM-endosomes (Fig. S1F, 3E). Means ± SEM, n≥3. p-value: ns; non-significant, *<0.05, **<0.01, ***<0.001 ****<0.0001.

MLC1 lacks N-linked complex glycosylation, which could have been used to separate the immature ER forms from the processed post-Golgi ones. Instead, we used the association of GlialCAM with MLC1^31, 34^ to selectively isolate the cell surface ubiquitinated MLC1. MLC1/GlialCAM was cell surface immunoprecipitated (cs-IP) with anti-GlialCAM antibody (Ab) bound to the cell surface. The affinity-purification followed by the denaturation and second IP for MLC1 was done to ensure examination of direct Ub-conjugation in MLC1 with anti-Ub Ab. Immunoblotting revealed increasing ubiquitination of mild P92S (∼5-fold) and severe C326R (∼40-fold) relatively to natively folded MLC1-wt at the cell surface (Fig. S1F).

As further evidence, we used MLC1-S280L with moderate turnover defect to determine the role of previously identified ubiquitin acceptor lysine (K) residues^38^ (Fig.1A). Mutagenesis of C-terminal K360 and cluster of lysines in the cytosolic loop (K172-175; 4K) decreased the PM ubiquitination (Fig.1C) and delayed the PM turnover of S280L (Fig.1D). Although the combination of these lysine mutations (5KA) further decreases Ub-conjugation, the S280L-5KA (T_1/2_ ∼2h) remained faster than MLC1-wt (Fig.1D, T_1/2_ >4h) indicating the involvement of additional factors for misfolded MLC1 faster endocytosis.

In summary (Fig.1E), comparison of PM turnovers after 1h chase for MLC1-wt (T_1/2_ >4h), P92S (T_1/2_ ∼1.5h), S280L (T_1/2_ ∼1h) and C326R (T_1/2_ ∼0.5 h)^34^ to their relative PM ubiquitination against native MLC1-wt (Fig.S1F, 3E) revealed a direct correlation between the faster PM turnover to the increased PM ubiquitination. Previous studies have also shown that the PM turnover of MLC1 variants is comparable in different cell types^31, 32, 34^.

### MLC1 mutations perturb cytosolic Ca^2+^ homeostasis and stimulate lysosomal exocytosis

Intracellular Ca^2+^ levels are potent regulators of proteostasis. MLC1 mutations cause endosomal compartment fusions without lysosomal permeabilization of protons^34^, which could relate to perturbations in the cytosolic Ca^2+^. The cytosolic Ca^2+^-transients upon activation of the store-operated Ca^2+^ influx from the extracellular space were examined using Fura-2 Ca^2+^-sensitive dye in MLC1-wt and S280L U251N cells^39^. In comparison to MLC1-wt, cells expressing MLC1-S280L exhibited diminished Ca^2+^-entry through the PM store-operated Ca^2+^-channels in response to intracellular Ca^2+^-stores depletion using thapsigargin (Tg) (Fig. 2A). Measurements of basal cytosolic Ca^2+^ levels using Calbryte520 and fluorescence reading showed a significant ∼2.5-fold increase in resting levels of cytosolic Ca^2+^ in S280L cells relative to MLC1-wt cells (Fig. 2B). The chronic intracellular Ca^2+^ increase implies dysregulation in the Ca^2+^ homeostasis, which can cause observed proteostasis stress to endosomal compartments. As a lysosomal Ca^2+^-response readout, we monitored the lysosomal Lamp1 exocytosis^40^ and hexosaminidase secretion^41^. The constitutive secretion of lysosomes in the absence of ionomycin and the inability of ionomycin (5μM) to stimulate lysosomal hexosaminidase exocytosis were consistent with the elevated cytosolic Ca^2+^ level imposing proteostasis stress to MLC1-P92S and S280L cells (Fig. 2C-D).

**Figure 2.**
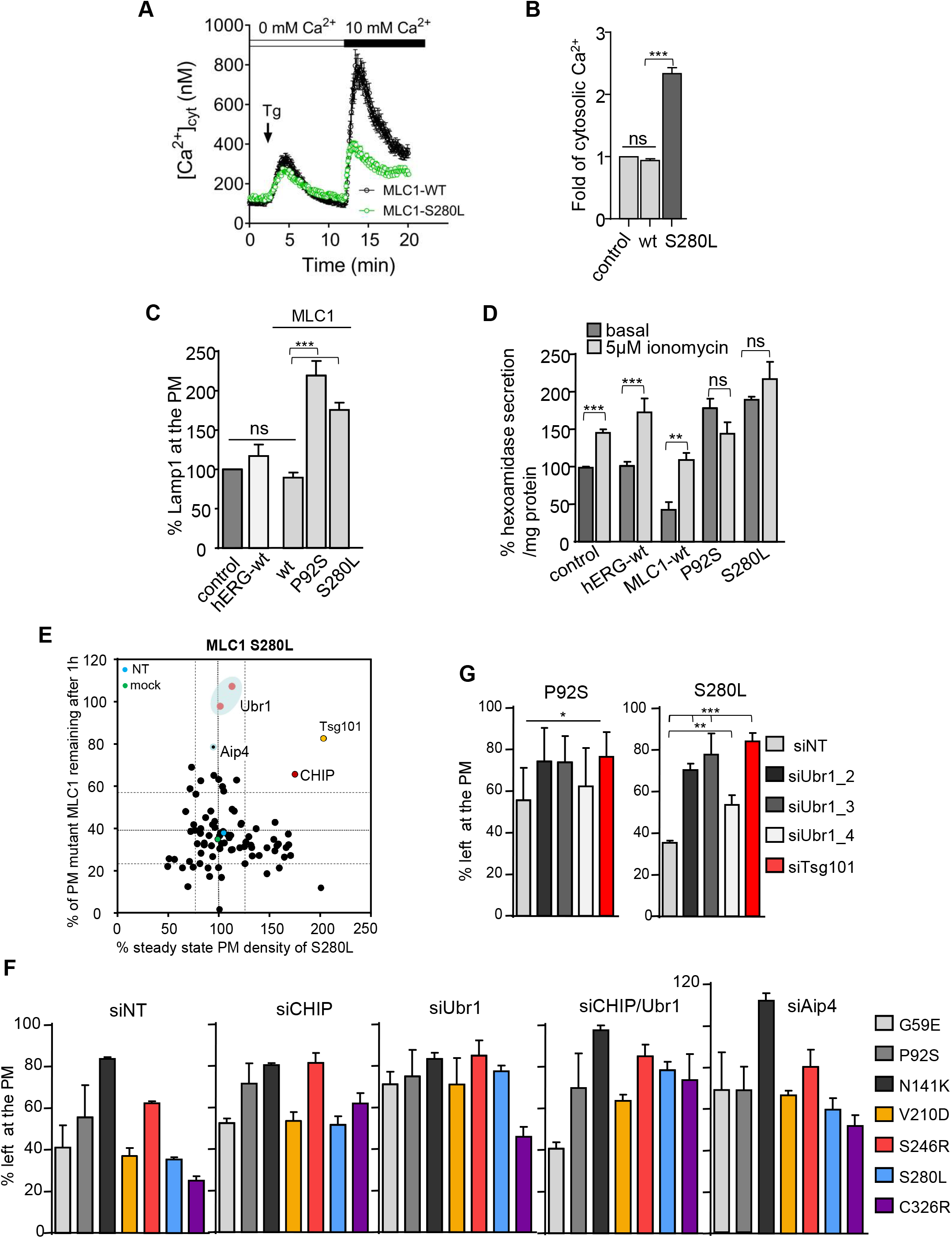
Screening post-Golgi and/or Ca^2+^-stress directed E3 ubiquitin QC-ligases. **A)** Comparison of activation of intracellular store-operated Ca^2+^ and extracellular Ca^2+^ entry channels across the PM (height of the peak) between HA-MLC1-wt and S280L expressing stable inducible U251N cells. Fura-2 was loaded in a Ca^2+^-free solution into live cells followed by thapsigargin (Tg) to induce an increase in the cytosolic [Ca^2+^]. The extracellular 10mM [Ca^2+^] was added to activate Ca^2+^ entry across the PM through Ca^2+^-permeable channels. n = 80-100 cells for each trace. **B)** MLC1 mutations cause basal cytosolic Ca^2+^ increase. Basal steady-state intracellular Ca^2+^-fold difference between control, MLC1-wt and S280L was measured using Calbryte520 in stable inducible U251N cells. Calbryte520 was loaded to live cells and measured using a fluorescence microplate reader and the signal was normalized to the number of cells. **C)** Basal Lamp1 expression at the PM indicating intracellular Ca^2+^-increase was measured using cs-ELISA (as in Fig.1B) in MLC1-wt, P92S, S280L stable inducible U251N cells and control and hERG channel expressing cells. **D**) Release of lysosomal enzyme β-hexosaminidase was measured using fluorometric assay from the medium of mock and 5μM ionomycin treated cells as in C. **E)** Phenotypic siRNA screen of candidate QC E3-ligases for the PM-endosomal clearance of MLC1-S280L causing intracellular Ca^2+^ overload. The PM expression and stability of HA-MLC1-S280L were measured using cs-ELISA (as in Fig1B) expressed in stable inducible HeLa. ESCRT-I protein Tsg101 (siTsg101) was used as a positive and siNT as a negative control. The potential PM/endocytic candidates CHIP, Aip4 and Ubr1 are indicated and lines of 3±SD. **F)** Cells were depleted with siRNA for QC E3-ligases CHIP, Ubr1, CHIP/Ubr1 and Aip4 and control siNT and the PM turnover of disease associated MLC1 variants^34^ were measured using cs-ELISA (as in Fig.1B). **G)** Ubr1 depletion decreases the PM turnover of misfolded MLC1. Cells were depleted using three different Ubr1 siRNA target sequences. siNT served as a negative and siTsg101 as a positive control. Cs-ELISA was used to measure the PM turnover after the 1h chase (as in Fig.1B). Means ± SEM, n≥3. p-value: ns; non-significant, *<0.05, **<0.01, ***<0.001.

### Identification of QC E3-ligases in post-Golgi compartments

The mechanism(s) of how endosomal compartments release proteostasis stress is not understood. To identify Ub E3-ligases that dispose of non-native endosomal membrane proteins and/or protect endosomal QC during Ca^2+^-perturbations, we performed a cell-based phenotypic E3-ligase siRNA sub-library screen against E3-ligases that may be involved in PQC. The objective was to select E3-ligases that impede the accelerated PM-endosomal turnover of MLC1-S280L that caused Ca^2+^ changes.

The PM turnover of MLC1-S280L was measured after one hour chase at 37°C using cs-ELISA in siRNA-transfected HeLa cells lacking endogenous MLC1. Targets that delayed the turnover more than three standard errors from the non-target siRNA (siNT) controls were considered hits (Fig. 2E). siRNA against TSG101, which is indispensable for ESCRT-dependent lysosomal degradation of ubiquitinated misfolded PM proteins, was used as a positive control^6, 7, 9, 42^.

We further examined found targets CHIP, AIP4 and Ubr1. CHIP is involved in PQC at multiple organelles^6, 7, 9^ and AIP4 (Itch) ubiquitinates cytosolic toxic misfolded proteins ^43^ and a subset of native endosomal receptors and adaptor proteins^44^. The n-recognin function of Ubr1 targets N-degrons in short-lived cytosolic and unfolded proteasomal substrates and outside its N-degron role, yeast ER membrane and mitochondrial proteins^17, 45–47^. As siAIP4 altered the endo-lysosomal transfer kinetics of constitutively ubiquitinated and recycling model cargoes (CD4t-Ub_n_, CD4cc-UbRΔG_4_, and the transferrin receptor (TfR)^48–50^ measured by cs-ELISA (Fig. S2C-D) and indicating functions outside QC, we focused on CHIP and Ubr1.

Measuring the PM-endosomal kinetics with cs-ELISA showed that depletion of Ubr1 (siUbr1) and CHIP (siCHIP) decreased the PM turnover of six transmembrane/cytosolic MLC1 disease mutations without an additive effect with siCHIP/siUbr1 or influence on the MLC1-wt or the native-like N141K mutation in the extracellular part of MLC1 (Fig. 2F, S2B). The specificity of siUbr1 to diminish the fast PM turnover of P92S and S280L was confirmed with three different siRNA targets (Fig.2G). As further evidence toward misfolded proteins, Ubr1 depletion did not influence constitute recycling of TfR or the lysosomal targeting of the Ub-dependent dependent (CD4t-Ub_n_, CD4cc-UbRΔG_4_) and Ub-independent (CD4t-LAMP) model transmembrane cargoes^48–50^ (Fig. S2C-D). These results suggest that CHIP and Ubr1 have PQC specificity toward misfolded MLC1 without targeting constitutive Ub-dependent cargo sorting by the ESCRT-machinery.

CHIP typically functions with chaperones/co-chaperones that enhance recognition of non-native conformations^51–53^. The cell surface IP (cs-IP) of MLC1/GlialCAM and immunoblotting revealed the highest ∼40-80-fold association of Hsc70, Hsp90, and Hsp40 (DNAJB1) with the severe C326R and half of that with milder P92S paralleling the severity in their PM defects. DNAJB1 type Hsp40 recruits misfolded substrates to Hsp70 as well as targets a range of human Hsp70-based disaggregase clients^54^. CHIP association increased ∼2-5-fold in mutants relative to that of MLC1-wt (Fig. 3A). The cs-ELISA assay showed that depletion of CHIP attenuated ∼66% of the PM turnover and ∼50% of endosomal internalization of the misfolded MLC1-P92S, which was reversed by CHIP-wt overexpression (Fig. S3A-C). The amount of chaperone/CHIP recruitment correlated with the severity of misfolded PM/endosomal MLC1 in the cluster, although misalignment of other members (e.g. in GlialCAM or ClC2) could contribute to recruitment^31, 32^.

**Figure 3.**
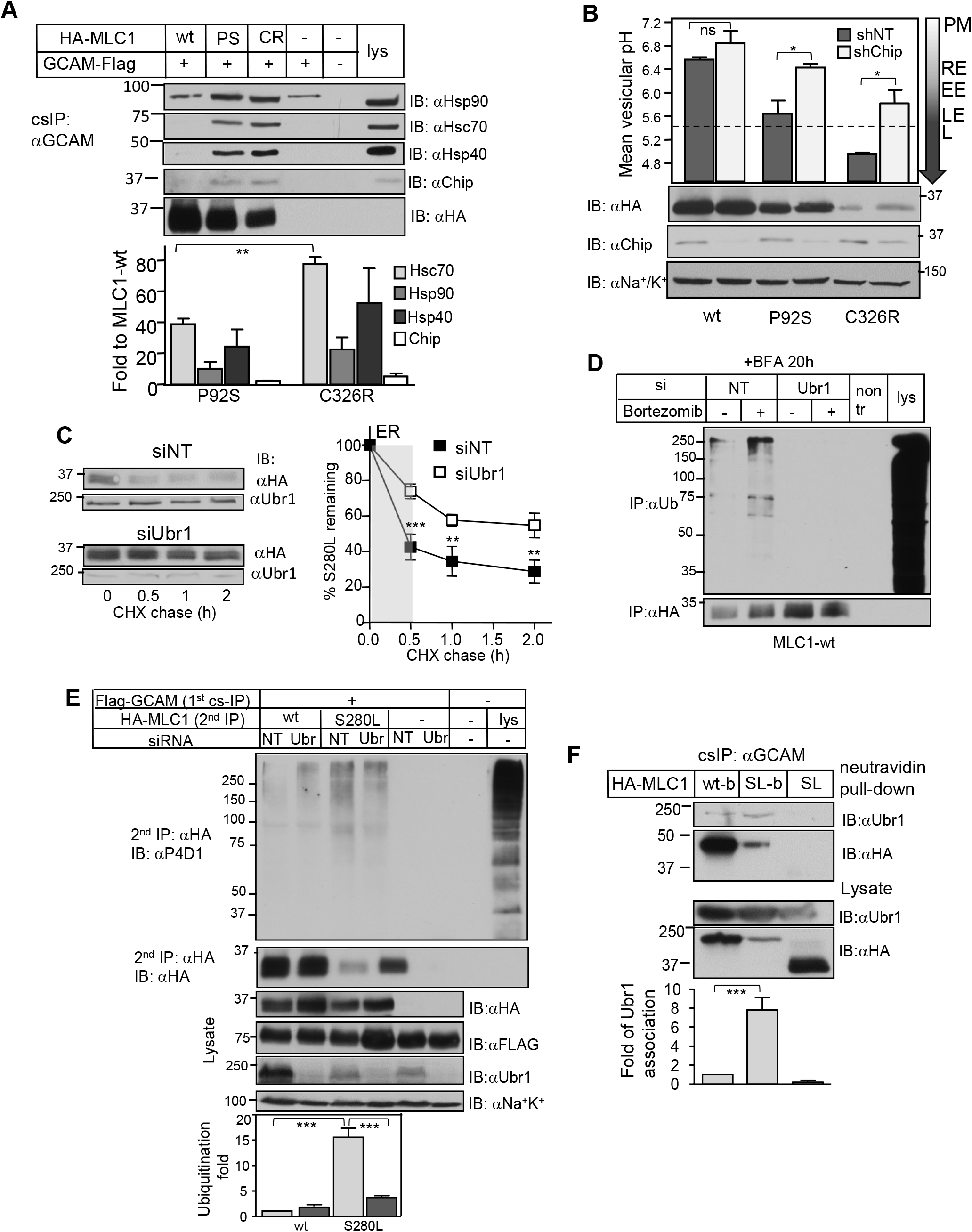
Disease-causing MLC1 variants are substrates for Ubr1. **A)** Interaction of molecular chaperones and E3 QC-ligase CHIP with misfolded MLC1 was monitored at the PM-endosomes. The MLC1/GlialCAM complex was cell surface immunoprecipitated (cs-IP) using GlialCAM Ab and protein interactions analyzed using Western blotting. Flag-GlialCAM was transiently expressed in stable inducible HA-MLC1 HeLa. The chaperone/CHIP amount was normalized to MLC1 expression at the PM. GCAM, GlialCAM; lys, lysate; α, anti. **B)** Mean single vesicular pH was measured from MLC1 containing endosomes using live-cell microscopy after 1h chase in cells depleted for CHIP with a short hairpin (sh) or shNT. The PM MLC1 was labelled with a pH-sensitive fluorophore and endosomal pH was measured as in Methods. The average weighted mean was calculated from at least three independent experiments, and >250 vesicles from ∼25-50 cells/ experiment were analyzed. Cells were as in A. Tha PM, recycling endosome (RE) pH 6.3-6.2, early endosome (EE) and late endosomes (LE) /lysosomes (L) <5.5 are indicated. **C)** The effect of Ubr1 depletion on MLC1-S280L total protein turnover was measured using CHX-chase and Western blot analysis. Control cells were treated with siNT. Cells were as in A. **D)** ERAD and Ubr1 were monitored in siRNA treated cells for Ubr1 or siNT. BFA was added for 20h to prohibit MLC1-wt export from the ER. Proteasomal degradation was inhibited with Bortezomib for 2h to accumulate ubiquitinated non-native MLC1. The ubiquitination signal was monitored after denaturation and IP for MLC1. Cells were as in A. **E)** The effect of Ubr1 depletion on PM-endosomal ubiquitination of native MLC1-wt and misfolded S280L was measured using cs-ubiquitination assay and quantification as in Fig. 1C. Cells were as in A. **F)** Proximity-dependent biotin protein-protein interaction assay between MLC1 and Ubr1 was probed in cells expressing transiently HA-MLC1-wt-Bir* (wt-b), S280L- Bir* (SL-b) or S280L (SL). Cells were treated with 50μM biotin for 20h and MLC1 was isolated using cs-IP (as in A). Biotinylated Ubr1 was recovered on neutravidin beads and analyzed using Western blotting. The biotinylated Ubr1 amount was normalized for the PM amount of MLC1. GCAM, GlialCAM; lys, lysate; α, anti. Means ± SEM, n≥3. p-value: ns; non-significant, *<0.05, **<0.01, ***<0.001.

If endosomal ubiquitination and ESCRT-dependent lysosomal targeting of non-native MLC1s are influenced by the dynamics of Ub-conjugation and deubiquitination^49, 55^, CHIP depletion should delay their endosomal transfer kinetics. To determine the direct endosomal kinetics of cs-labelled and endocytosed MLC1 variants, we used pH-sensitive single vesicle fluorescence microscopic analysis in live cells. MLC1 variants endosomal location was based on the progressive acidification of the endo-lysosomal vesicular pH (pH_v_), ranging from pH ∼7.2 to 4.5^6, 7, 56–58^. MLC1-wt was confined to recycling endosomes (pH_v_ 6.3±0.2) after endocytosis at 37°C for 1 h, while the P92S (pH_v_ 5.65±0.3) and C326R (pH_v_ 5.0±0.04) were targeted to the MVB (late endosomes)/lysosomes based on the characteristic endosomal pH_v_ (Fig.3B). The mutant delivery to MVB/lysosomes was delayed by depletion of CHIP or ESCRT-0 and -I constituents STAM, Hrs or TSG101 (Fig. 3B and S3D-E). These data indicate that the accelerated PM/endosomal clearance of a fraction of misfolded MLC1 is mediated by the ubiquitination- and ESCRT-dependent PQC.

### Ubr1-dependent ubiquitination of MLC1 variants

Lack of CHIP caused only partial inhibition of the endo-lysosomal transfer of MLC1 variants (Fig. 3B). The found second E3-ligase Ubr1 has been suggested to have PQC functions outside its role as n-recognin, mostly in yeast, toward a limited number of cytosolic, misfolded ER, mitochondrial and cytosolic PQC clients^45–47^. Ubr1 in the human ER or endosomal PQC is not well understood. We selected MLC1-S280L with moderate expression and turnover defect (Fig. 1A, 1E) for further QC studies.

First, to address the cellular and the ER role of Ubr1, the disappearance kinetics of MLC1-S280L was measured upon translational inhibition by cycloheximide (CHX) and immunoblotting in siUbr1 and siNT treated cells. The fast degradation rate of MLC1-S280L during the first 0.5 h chase was inhibited by ∼50% in siUbr1 treated cells (Fig. 3C), suggesting that Ubr1 contributes to the ER clearance of misfolded MLC1. To address the ER effect of Ubr1, non-native MLC-wt was accumulated in the ER by preventing its ER exit with Brefeldin A (BFA) and ERAD with proteasomal inhibitor Bortezomib. The ubiquitination of non-native MLC-wt was reduced by >75% in siUbr1 treated cells (Fig. 3D). These results showed that a fraction of unassembled or incompletely folded MLC1 is targeted for Ubr1-dependent ERAD.

Next, we assessed Ubr1 capacity for Ub-conjugation at the PM-endosomes. MLC1/GlialCAM was isolated using cs-IP followed by a second IP for MLC1 after denaturation to reveal direct MLC1 ubiquitination using immunoblotting and anti-Ub Ab (Fig. 3E, as in Fig. 1C, S1F). Ubiquitination was normalized to the MLC1 amount. Ubr1-depletion decreased the ubiquitination of misfolded MLC1-S280L by ∼75% at the PM, while its PM amount increased ∼3-fold (Fig. 3E, *lower panel*) without affecting natively folded MLC1-wt at the PM.

To demonstrate the interaction of Ubr1 with MLC1 at the PM-endosomes, misfolded MLC1-S280L was expressed with Flag-Ubr1-wt or catalytically inactive (Ubr1-CI). After mild in vivo cross-linking, S280L/GlialCAM was isolated using cs-IP without denaturation. Ubr1- wt reduced and Ubr1-CI elevated S280L amount at the PM, however, Ubr1 detection was low. As a more sensitive in vivo approach, we measured the MLC1/Ubr1 interaction with the proximity biotinylation technique using MLC1-Bir* as the bait and cs-IP^59^. The endogenous Ubr1 displayed ∼8-fold higher biotinylation susceptibility by the MLC1-S280L-Bir* than MLC1-wt (Fig. 3F). These data showed that the non-native astrocytic regulatory cluster membrane protein MLC1 variant represents previously unrecognized PQC clients for Ubr1 in the ER and the PM-endosomes.

### Ubr1 functions as an endosomal E3 ubiquitin QC-ligase

Ubr1 subcellular location in human cells is not well known and was evaluated using indirect immunostaining and fluorescence microscopy for endogenous-Ubr1, Flag-Ubr1 and mCherry-Ubr1 without the presence of misfolded protein or Ca^2+^-stress. The majority of Ubr1 was associated with membrane-puncta and partly with the PM (Fig. S4A), phalloidin stained F-actin at endosomal invaginations, as well as the clathrin adaptor (AP2) and the clathrin light chain (CLC) (Fig. S4B) being consistent with its role at the PM-endosomal compartment. Additionally, Ubr1 was found partly in the nuclei (Fig. S4A) and with the ER-markers KDEL and ERp57, but not with the lysosome marker Lamp1 or autophagosome marker LC3b (Fig. S4B).

Next, the Flag-Ubr1 colocalization was assessed in the presence of endocytoses misfolded MLC1-S280L and native MLC1-wt. Extracellular MLC1 HA-epitope was selectively labelled at the PM and endocytosed 30 min for immunofluorescence or proximity ligation assay (PLA)^31, 32^ to visualize protein-protein interactions. Both Ubr1 and MLC1-S280L colocalized with early endosomal marker EEA1 (Fig. 4A-B). We confirmed the preferential interaction of endosomal misfolded MLC1-S280L with the Ubr1-wt and Ubr1-CI, as well as EEA1+ using the PLA (Fig. 4C-D). The immunolocalization and proximity interaction were substantially enhanced between Ubr1-CI and S280L over the MLC1-wt at the PM proximity (Fig. S4C), reinforcing the notion that Ubr1 interacts with misfolded MLC1 specifically at PM-endosomal compartments (Fig. 3E).

**Figure 4.**
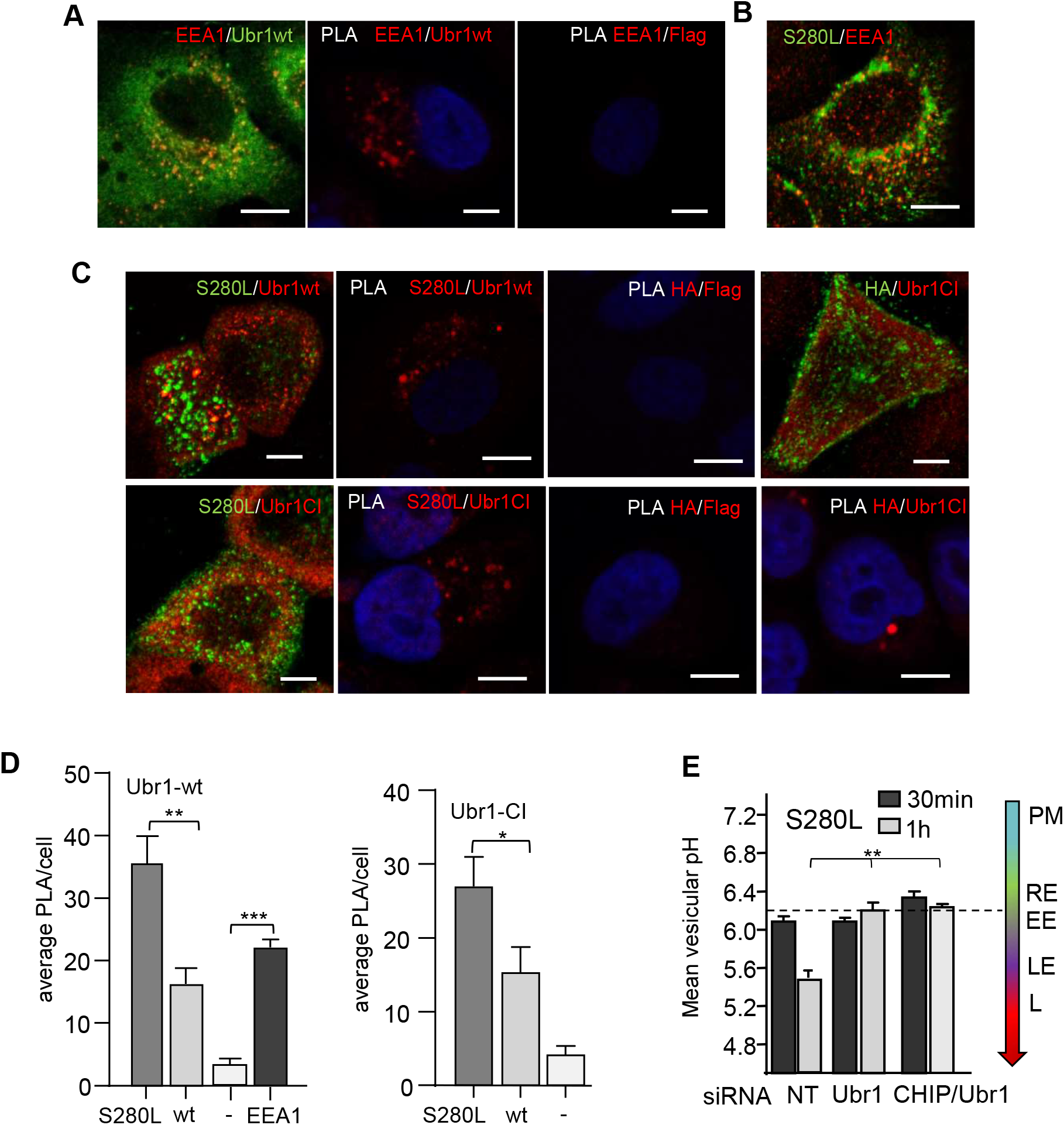
Ubr1 functions as an endosomal QC E3 ubiquitin ligase. **A)** Immunofluorescence microscopy for early endosomal antigen EEA1 and Ubr1 interaction was detected using anti-EEA1 and anti-Flag Abs, and protein-protein interaction with PLA was performed between EEA1 and Flag epitopes. Cells without Flag-Ubr1-wt expression were used as a control. Bar 5 μm. **B)** Immunofluorescence colocalization of internalized (30min) endosomal MLC1-S280L was visualized using anti-HA with early endosomal EEA1. **C)** Immunofluorescence of endosomal MLC1-S280L or MLC1-wt and Ubr1 interaction was determined for internalized anti-HA after 30min and stained for Flag-Ubr1-wt or CI using anti-Flag. PLA was performed between HA-Flag epitopes and cells without MLC1 were used as a control. Bar 5 μm. **D)** Quantification of average protein interaction with PLA between Flag-Ubr1-wt or CI and MLC1-wt, S280L or EEA1 from>30 cells. **E)** Mean vesicular pH measurement of S280L containing endosomes was determined using live-cell single fluorescence microscopy in cells depleted for CHIP, Ubr1, CHIP/Ubr1 or control siNT. PM; plasma membrane, RE; recycling endosome (pH 6.3-6.2), EE; early endosomes, LE; late endosome, L; lysosome. Means ± SEM, p-value: *p<0.05, **p<0.01, ***<0.001.

Next, the significance of Ubr1 on the endosomal sorting of MLC1 was interrogated by monitoring misfolded MLC1-S280L endo-lysosomal trafficking with pH_v_-analysis (as in Fig. 3B) in control and siUbr1-depleted cells after 0.5 and 1 h chase. Ubr1 depletion prevented the delivery of MLC1-S280L from early endosomes (pH_v_ 6.1±0.08) to late-endosome/MVB (pH_v_ 5.5±0.02) during the 30-60 min chase period (Fig. 4E). A comparable effect was documented upon CHIP and Ubr1 depletion (Fig. 4E). Ubr1 depletion did not affect fluid-phase endocytosis, probed by dextran uptake or the lysosomal targeting of a ubiquitinated model substrate CD4t-Ub_n_ (Fig. S4D, see also Fig. S2C-E), ruling out its non-specific effect on ESCRT-dependent ubiquitinated cargo sorting. Jointly, these results strongly suggest that Ubr1 activity is indispensable for the timely delivery of misfolded endosomal MLC1 variants into MVB/lysosomes.

### Ubr1 is activated by proteostasis stress in endosomal compartments

The role of Ubr1 in a cytosolic and ER QC has been demonstrated upon cellular stress in yeast^46, 60^. We sought evidence for Ubr1 connection to endosomal proteostasis stress in human diseases using MLC1 mutant cells with endosomal fusions^34^ and perturbed Ca^2+.^-homeostasis (Fig.2A-D). MLC1-pool was selectively labelled with anti-HA and endocytosed (30 min, 37°C), and compared to Lamp1+ lysosomes using immunofluorescence microscopy. Only MLC1-S280L but not wt provoked the accumulation of enlarged Lamp1+ late endosomes/lysosomes (Fig.5A). Importantly, this phenomenon was suppressed by Ubr1 depletion with siUbr1 (Fig.5A and C-D) suggesting that Ubr1 activity/binding with misfolded endosomal MLC1 is responsible for the late endosome/lysosome enlargement (Fig.5A-D).

Remarkably, enlarged Lamp1+ late endosomes/lysosomes were positive for LC3b and MLC1-S280L identifying them as autolysosomes (Fig.5B-D). A subpopulation of LC3 and MLC1-S280L positive vesicles, lacking Lamp1 staining, likely represent newly formed cargo-selective autophagosomes (or endo-phagosomes) (Fig. 5B).

**Figure 5.**
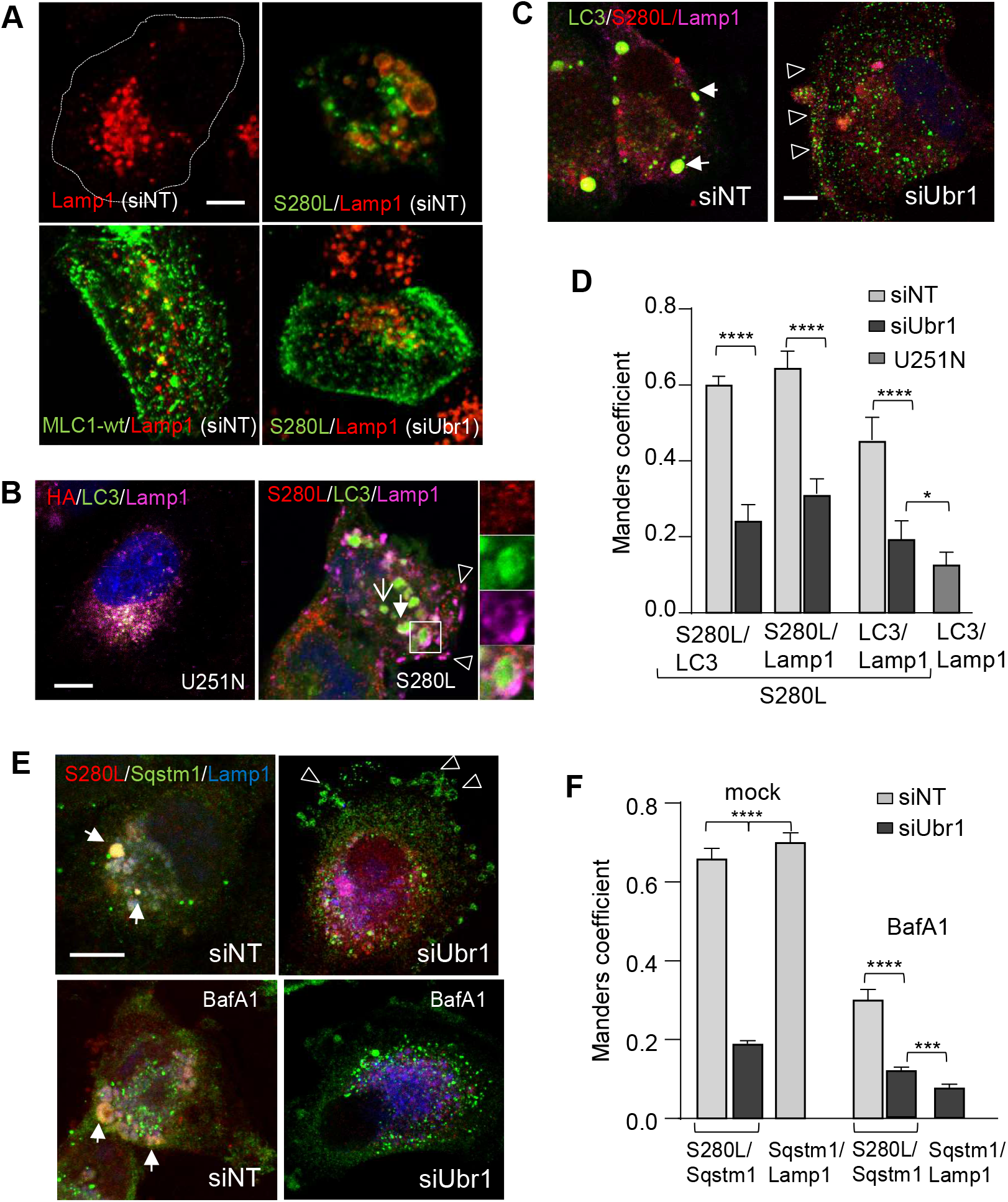
Ubr1 is required for the endo-phagy formation. **A)** Immunofluorescence microscopy of the Ubr1 expression effect on the endo-lysosomal pathway morphology was monitored in siRNA treated cells for Ubr1 or control NT. Endosomal MLC1-S280L or wt was internalized (30min) and visualized using anti-HA Ab and Lamp1 was used as a lysosomal marker to detect the enlarged phenotype. Bar 5 μm. **B)** Autophagosome marker LC3b and lysosomal marker Lamp1 were used to identify autolysosomes (arrow, insert) and autophagosomes (open arrow) in MLC1-S280L cells as in A. Arrowhead points to exocytosed lysosomes at the PM. Bar 5 μm. **C)** Endosomal MLC1-S280L and Ubr1 connection to the endosomal stress and formation of autophagosomes/autolysosomes (endo-phagy) was monitored in Ubr1 depleted (siUbr1) or control siNT cells as in A. LC3b was used as an autophagosome and Lamp1 as a lysosomal marker. The arrow indicates autolysosomes positive for S280L, Lamp1 and LC3b, and the arrowhead points to the PM relocated S280L, Lamp1 and LC3b. Bar 5 μm. **D)** Manders’ overlap coefficient of indicated autophagy (LC3b) and lysosome (Lamp1) markers. Representative cells are in B-C. n>40 cells. **E)** Connection of Ubr1 and endosomal MLC1-S280L to the endosomal stress and recruitment of autophagy scaffold SQSTM1/p62 to phagophores in MLC1-S280L cells was done as in A. BafA1 was used to inhibit autophagosome-lysosome fusions, which resulted in the cytosolic dispersion of SQSTM1/p62. The arrow indicates autolysosomes and arrowheads the SQSTM1/p62 positive PM blebs. Bar 5 μm. **F)** Manders’ overlap coefficient of selective autophagy scaffold SQSTM1/p62 and lysosomes (Lamp1) with HA-MLC1-S280L. Representative cells are in E. n>26 cells. Means ± SEM, p-value: *p<0.05, ***p<0.001, ****p<0.0001.

Interaction of Ubr1 with misfolded endosomal MLC1 and the formation of large Lamp1/LC3b positive autophagosomes/autolysosomes suggest that Ubr1 has endo-lysosomal stress QC functions. As support, Ubr1-depletion by siRNA abrogated the formation of autolysosomes and their precursors (endosome-derived autophagosome) dispersing LC3b to the cytosol and PM resulting in ∼ 3-fold decrease in colocalization of S280L/LC3b and S280L/Lamp1 (Fig. 5C, D). Importantly, formed autolysosomes were positive (SQSTM1/Lamp1) for the stress-induced SQSTM1/P62 scaffold that targets ubiquitinated proteins for selective autophagy^61^ (Fig. 5E-F). Consistently, lac of Ubr1 prevented the accumulation of SQSTM1/P62 into the forming autophagosome, dispersed SQSTM1/P62 into the cytosol and endosomal anti-HA labelled MLC1-S280L to low-curvature PM blebs. Ubr1 depletion decreased S280L/SQSTM1 colocalization by ∼3-fold (Fig. 5E, also 6E).

Inhibiting the autophagosome-lysosome fusion by bafilomycin A1 (BafA1) resulted in cytosolic dispersion of SQSTM1/p62 and the reduced colocalization of the SQSTM1/p62 with MLC1-S280L, consistent with the endosomal mutant degradation by auto-lysosomal pathway (Fig. 5E-F). Furthermore, Western blot analysis revealed that the cellular SQSTM1/p62 and the autophagy membrane LC3b-II/I ratio were increased upon siRNA-mediated Ubr1 depletion in MLC1-S280L cells as well as in control cells (Fig. 6A-B). This suggests that Ubr1 has a substantial role in general proteostasis QC. Consistently, the lack of Ubr1 resulted in the accumulation of pre-autophagic isolation membranes with compromised maturation to autophagosomes pinpointing its importance for selective autophagy (Fig. 5C-F, 6E).

**Figure 6.**
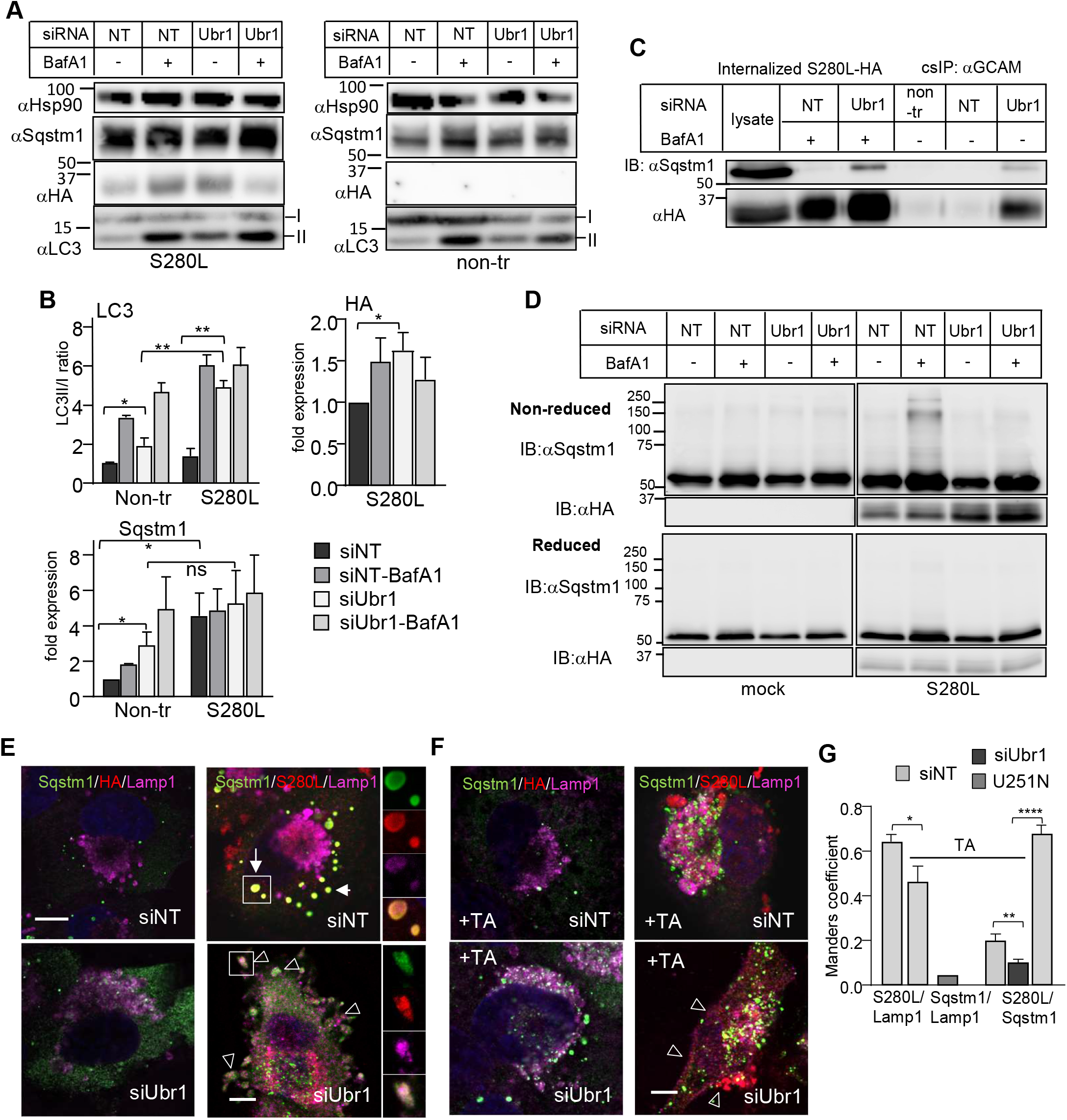
Ubr1 is a proteostasis stress QC-ligase for the selective SQSTM1/p62 endo-phagy pathway. **A)** Western blot analysis of cells depleted with siRNA for Ubr1 or NT to measure the expression of SQSTM1/p62 and the formation of mature autophagosomes using lipidated/cytosolic LC3b-II/I ratio in HA-MLC1-S280L and non-expressing U251N cells. BafA1 was used to inhibit autophagosome-lysosome fusions to decrease autophagosome-cargo degradation. Hsp90 was a loading control. **B)** Densitometric quantification of MLC1-S280L, SQSTM1/p62 and lipidated/cytosolic LC3b-II/I ratio from A. n=3. **C)** Protein-protein interaction between autophagy scaffold SQSTM1/p62 and endosomal MLC1-S280L was done using cs-IP (as in Fig.3A) for cells treated as in A. **D)** Oligomerization of SQSTM1/p62 requires Ubr1 in the presence of MLC1-S280L. Cells treated as in A were analyzed in reducing and non-reducing conditions and Western blot analysis. **E-F)** Immunofluorescence microscopy of Ubr1 and arginylation effect on SQSTM1/p62 recruitment and the endo-lysosomal pathway morphology was monitored in HA-MLC1-S280L or non-expressing U251N cells treated with siRNA for Ubr1 or control NT. Endosomal MLC1-S280L and control were internalized (30min) and visualized using anti-HA Ab, and colocalization with Lamp1 lysosomal marker and SQSTM1/p62 selective autophagy scaffold protein (as in 5E) was determined +/− of arginylation inhibitor, tannic acid. The arrow indicates autolysosomes and arrowheads lysosomal Lamp1 or SQSTM1/p62 at the PM and in blebs. The insert illustrates S280L containing autophagosome (upper panel) or the PM trapped cargo phagophore (lower panel). Bar 5 μm. **G)** Manders’ overlap coefficient of selective autophagy scaffold SQSTM1/p62 and lysosomes (Lamp1) with HA-MLC1-S280L. Representative cells are in E-F. n>29. non-tr, non-transfected; α, anti; Means ± SEM, p-values: *p<0.05, **p<0.01, ****p<0.0001.

Notably, the PM-endosomal MLC1-S280L/GlialCAM isolated using cs-IP (as in Fig.3A) showed that SQSTM1/p62 interacted with the S280L/GlialCAM without Ubr1 and in BafA1 treated cells (Fig. 6C). Importantly, this fractional interaction of SQSTM1/p62 (and LC3b) without Ubr1 to endosomal S280L indicated an alternative binding mechanism than Ub/Ubr1.

Instead, Ubr1 was required for the maturation of autophagosomes (Figs 5C-F). The oligomerization of multivalent SQSTM1/p62 allows the simultaneous selection of ubiquitinated cargo and pre-autophagic isolation membranes during selective autophagy^62^. Accordingly, the inhibition of the misfolded endosomal MLC1-S280L transfer to lysosomes with BafA1 led to the disulphide-dependent SQSTM1/p62 oligomerization^63^ as oligomers were eliminated under reducing conditions (Fig. 6D). Notably, the lack of Ubr1 completely abrogated the oligomerization of SQSTM1/p62 (Fig. 6D).

Collectively, the data so far (Fig.1–6D) indicated that misfolded MLC1 changes intracellular Ca^2+^ homeostasis and renders proteostasis stress to the endosomal pathway culminating in the activation of Ubr1- and SQSTM1/p62-dependent cargo selective endo-phagy.

### Ubiquitination and arginylation cooperatively safeguard endo-lysosomal proteostasis

SQSTM1/p62 was able to bind misfolded endosomal MLC1 without Ubr1 (Fig. 5E, 6D) implying alternative cargo recognition. SQSTM1/p62 was recently found to bind N-degrons, including arginine (R), that facilitates disulphide bond-linked aggregation of SQSTM1/p62 and SQSTM1/p62 interaction with LC3, as well as its cargo delivery to the autophagosome^19^. Arginylation can occur on exposed N-terminal residues or internally located side-chains of proteins^64^. To assess whether SQSTM1/p62 can recognize misfolded MLC1 via arginylation, arginyltransferase was inhibited by tannic acid in MLC1-S280L cells and analyzed using immunofluorescence microscopy. Tannic acid induced the SQSTM1/p62 dispersion prohibitin autophagosome formation in S280L cells similar to lack of Ubr1 (Fig. 6F-G, S5A). Inhibition of arginylation and Ubr1 depletion prevented the formation of autophagosomes and enhanced S280L and Lamp1 accumulation at the PM (Fig.6F-G, S5A).

Arginylated glutamic (E) or aspartic acid (D) can be recognized by the arginylation specific antibody. Arginylation colocalized with Ubr1 to membrane puncta and autolysosome-like structures in S280L cells (Fig. 7A, S5B). In Ubr1-depleted S280L cells, argynylated substrates preferentially accumulated at the PM with LC3b (Fig. 7B, S5B).

**Figure 7.**
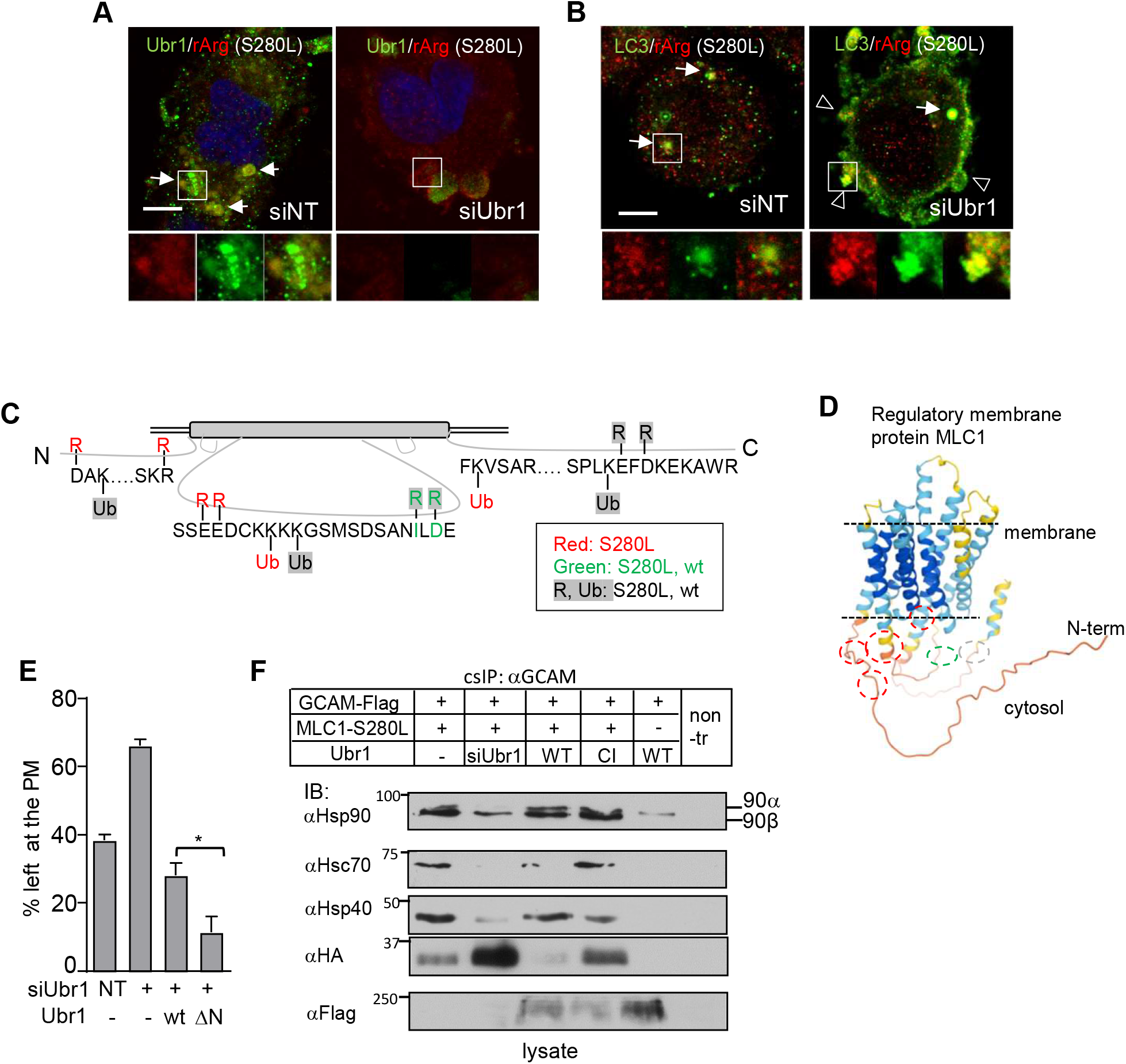
Proximity clustering of ubiquitin/arginine and Ubr1-Hsp90 chaperone complex mediate cargo selective endo-phagy. **A)** Immunofluorescence microscopy of Ubr1 and arginylated substrate colocalization to membrane puncta and autolysosome-like structures in MLC1-S280L expressing U251N cells depleted for Ubr1 or NT by siRNA. The insert illustrates the individual staining pattern of the Ubr1/rArg containing structure. Bar 10 μm. Manders’ overlap coefficients are in Figure S6B. **B)** Immunofluorescence microscopy of autophagosome marker LC3b and arginylated substrate (rArg) colocalization in MLC1-S280L cells as in A. The insert illustrates the individual staining pattern of LC3b/rArg. Bar 10 μm. Manders’ overlap coefficients are in Figure S6B. **C)** Mass spectrometry analysis of affinity-purified MLC1-wt and S280L amino acids at the intracellular part for arginylation and ubiquitination. Analyzed peptides are in Figure S5C. **D)** Ubiquitin/arginine residues from C indicated structural proximity clustering of ubiquitin/arginine residues in misfolded MLC1 (red circles). Red circles: R/Ub S280L, green: R wt/S280L and grey: R/Ub wt/S280l as in C. **E)** The UBR-box deleted Flag-Ubr1-ΔN and Flag-Ubr1-wt effect on the PM stability of MLC1-S280L was measured using cs-ELISA. Cells were depleted for Ubr1 or NT using siRNA and Flag-Ubr1 was transiently expressed. **E)** The effect of Ubr1 depletion and Flag-Ubr1-wt or CI expression on Hsp90β/Hsc70/Hsp40-complex association with misfolded MLC1-S280L at the PM-endosomes. Cells were prepared as in E and the PM-endosomal MLC1 was isolated using cs-IP and Western blot analysis as in Fig.3A. α, anti; GCAM, GlialCAM. Means ± SEM, p-values: *p<0.05, **p<0.01, ****p<0.0001.

The above data indicates that both arginylation and Ubr1-induced ubiquitination of misfolded MLC1 is required for the SQSTM1/p62-driven endo-phagosome formation. To test this, we monitored the arginylation and ubiquitination of the affinity-purified MLC1-wt and S280L by tandem mass spectrometry (Fig. 7C, peptides S5C). Arginylation was found D31 and R42 residues in the N-terminal tail and E168-169 residues in the 2^nd^ cytosolic loop of S280L, adjacent to the Ub-acceptor sites (K33, K173 and 175) forming structural proximity clusters (Fig.7D, red circles). We also observed folding-independent arginylation in the wt and S280L (Fig. 7C, 7D green circle). Ubiquitin in K173 and 175 in the 2^nd^ cytosolic loop and with the C-terminal K360 (Fig.1A) was also identified with MS/MS, confirming MLC1 ubiquitination results in Fig.1C-D.

A recent study showed that proteostasis stress increased the cytosolic accumulation of R-BiP ^19^. Immunofluorescence microscopy confirmed that the R-BiP level was increased in the cytosol of S280L cells and R-Bip was also confined with LC3b, Lamp1 and S280L to autolysosomes (Fig. S6A-D). Considering that the recruitment of Ubr1 by arginylation to misfolded MLC1 was limited (Fig. 6E-F, S6A-D), we tested the effect of the N-terminal UBR-box deletion (ΔN), responsible for the recognition of N-terminal arginylation by Ubr1. The PM turnover of MLC2-S280L, measured by cs-ELISA, showed that ΔN-Ubr1 overexpression accelerated the PM turnover of S280L more than Ubr1-wt, suggesting that Ubr1 is recruited via an alternative mechanism (Fig. 7E). This is consistent with the Hsp70-dependent and N-degron-independent PQC function of Ubr1 in the cytosol^65^.

To evaluate the involvement of molecular chaperones in MLC1-S280L recognition by Ubr1, the PM resident MLC1 was isolated with cs-IP (as in Fig.3A) or cellular pool using IP from siUbr1-depleted, and Flag-Ubr1-wt or Ubr1-CI expressing cells followed by immunoblotting of interacting chaperones. Hsp90β (83.2kDa) preferentially interacted at the PM, while both Hsp90α (90kDa) and Hsp90β were interacting with the cellular MLC1-S280L (Fig. 7F, S6E). Lack of Ubr1 decreased Hsc70/Hsp40/Hsp90β association with the mutant in both compartments.

In summary, these results show that Ubr1 recruitment to misfolded endosomal MLC1 requires endosomal stress and an Hsp90-chaperone complex. Ubr1-client ubiquitination/arginylation drives SQSTM1/p62 oligomerization ensuring cargo selective endo-phagy maturation.

### Ubr1/ SQSTM1/p62 clear ubiquitin/arginine PQC clients during Ca^2+^ -induced stress

As a finale, we asked whether Ubr1 regulates a wider biological endosomal pathway stress QC. Because of the lack of markers for the endo-lysosomal stress, we used a morphological readout of Lamp1+ lysosomes (Fig. 5) in Hela cells.

First, as Ubr1 was found in endosomal stress QC, then the lack of Ubr1 should cause changes in lysosomes. Abrogation of Ubr1 (Fig.8A condition) by siRNA was sufficient to enhance the accumulation of arginylated proteins to lysosomes (Fig. 8B, E) and enlarge lysosomes (Fig.8B, F). Importantly, Ubr1 depletion with inhibition of arginyltransferase (TA) provoked pronounced enlargement of lysosomes, indicating that Ubr1 with arginylation are important for endosomal pathway health (Fig.8B, F).

**Figure 8.**
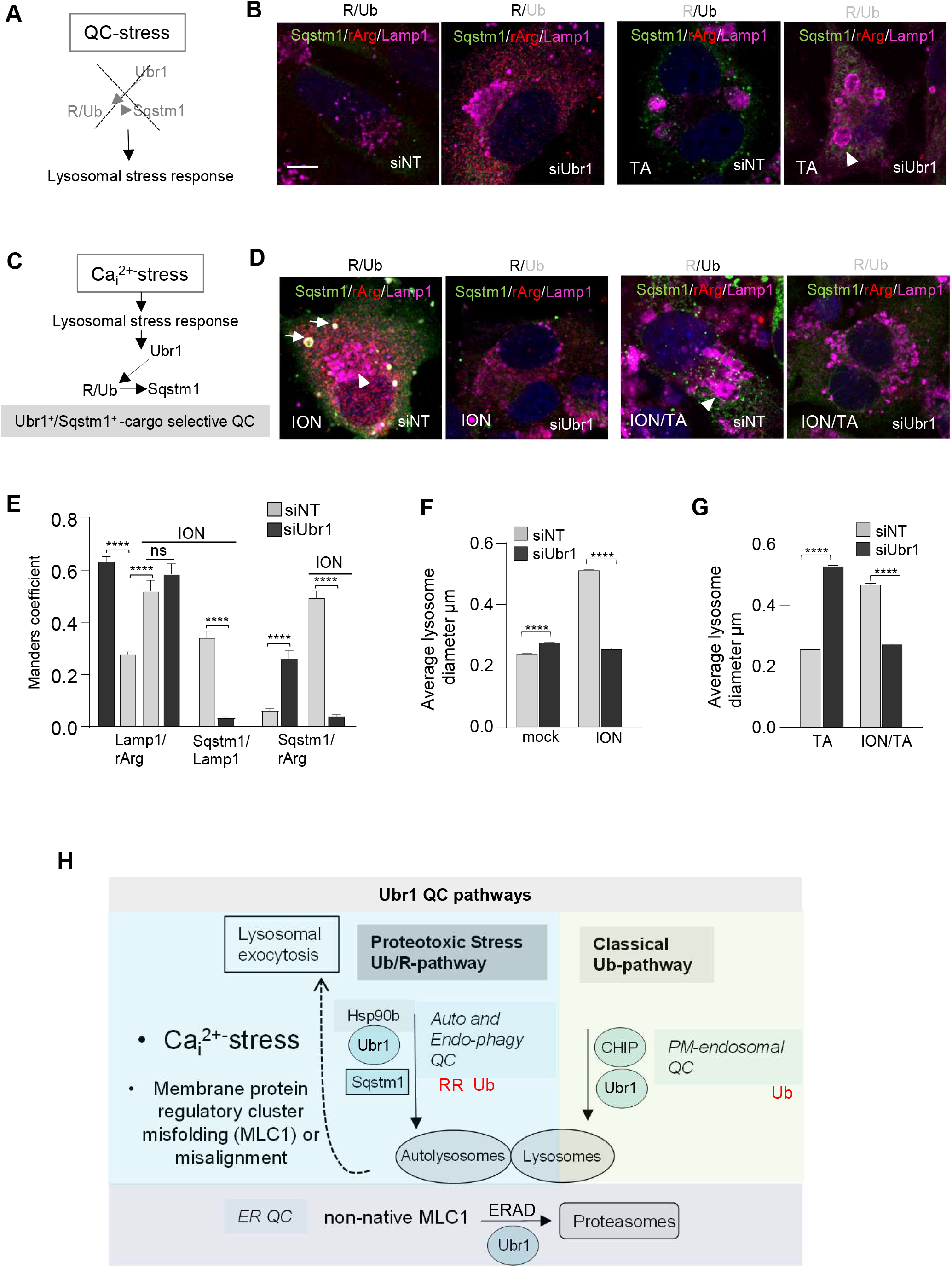
The interplay of ubiquitination and arginylation enhance Ubr1 and SQSTM1/p62-mediated Stress QC to safeguard endo-lysosomal pathway proteostasis. **A)** Diagram of the proposed role of lack of Ubr1 and arginylation signal resulting in QC-stress. **B)** Ubr1-dependent QC-stress. The morphological analysis of lysosomal response upon Ubr1 depletion with siRNA and the relation of arginylation signal and SQSTM1/p62 recruitment to lysosomes was determined using immunofluorescence microscopy in Hela cells. Lamp1 was a lysosomal marker and tannic acid (TA) was used to inhibit arginylation. The arrowhead indicates enlarged lysosomes. Bar 5 μm. **C)** Diagram of Ca^2+^ -stress leading to the proposed role of Ubr1 and SQSTM1/p62 in cargo selective QC. **D)** Ubr1-dependent Ca^2+^-stress QC. As in B, but ionomycin (ION) was used to induce intracellular Ca^2+^-stress. The arrow indicates autolysosomes and arrowhead enlarged lysosomes. Bar 5 μm. **E)** Manders’ overlap coefficient of autophagy scaffold SQSTM1/p62, arginylated substrates (rArg) and lysosomes (Lamp1) in siNT or siUbr1 treated HeLa cells and upon ionomycin Ca^2+^-stress (ION). Representative cells are in B and D. n>48 cells. **F-G)** Average lysosomal diameter from experimental conditions presented in B and D. n>2000 lysosomes. Means ± SEM, p-values: ****p<0.0001. **H)** Summary of proposed classical ubiquitin (Ub)-mediated QC and ubiquitin/arginine (Ub/R)-mediated Stress and Endo-phagy QC pathways. Ubr1 was found to have a previously unrecognized role as a new Endosomal and Stress QC-ligase at the endosomal autophagy (endo-phagy) pathway to support PQC during proteostasis stress and disease. Classical ubiquitin (Ub)-mediated QC pathway function toward lysosomes, which uses QC-ligase CHIP, ubiquitin-mediated endosomal cargo concentrating ESCRT machinery and endosomal Ubr1. However, proteostasis stress initiated either by intracellular Ca^2+^ overload or misalignment of the astrocyte regulatory MLC1 signalling cluster cause fused endosomal compartments with altered cargo sorting. This leads to the activation of an alternative Ub/R (arginine)-mediated Auto/Endo-phagy QC pathway, where Ubr1 drives ubiquitinated/arginylated clients to autophagy scaffold SQSTM1/P62 to enhance the maturation of cargo selective autophagosomes. Subsequently, Ubr1 and SQSTM1/p62 function as a rerouting surveillance mechanism to safeguard proteostasis at the endosomal compartments.

Ubr1 is important for endo-phagy stress QC caused by MLC1 mutants, which also elevated cytosolic Ca^2+^. Next, we tested the Ubr1 link to Ca^2+^-stress and whether it can clear a variety of clients via cargo selective SQSTM1/p62 autophagy (Fig.8C condition). Intracellular Ca^2+^ was elevated by Ca^2+^ ionophore, ionomycin, known to trigger proteostasis stress via mitochondrial oxidative stress^66^, the release of lysosomal enzymes (Fig. 2D), by reducing PtdIns(4,5)P2 level critical for the PM protein stability and endocytosis^67^ and imposing ER-stress^68^. Ionomycin-induced (ION) enlargement of Lamp1+ lysosomes, the appearance of Lamp1/SQSTM1/p62 positive autolysosomes and arginylation signal to lysosomes (Lamp1/rArg) with SQSTM1/p62 (Fig.8D, E). Lack of Ubr1 during Ca^2+^-stress abrogated SQSTM1/p62 expression, autolysosomes and decreased significantly lysosomal diameter (Fig.8D, F). Inhibition of arginylation abrogated selective autolysosome formation and partly the lysosomal enlargement being full with the removal of Ubr1 in Ca^2+^-stress (Fig.8D, F-G).

In summary, the PQC function of Ubr1 targets clients for SQSTM1/p62 to fuel ubiquitinated/arginylated cargo selective endo-phagy and autophagy during Ca^2+^-stress (Fig. S6F).

## Discussion

Here, we describe previously unrecognized actions of Ubr1 and SQSTM1/p62 in cargo selective endo-phagy/autophagy clearing specifically ubiquitinated/arginylated clients during proteostasis stress. We found Ubr1 important for attenuating proteostasis stress at the fused endosomal compartments with altered cargo sorting that were initiated either by conformationally challenged astrocyte signalling cluster MLC1 variants possessing proximity clustering of ubiquitin-arginine posttranslational modifications and/or cytoplasmic Ca^2+^-overload. In turn, the lack of Ubr1 alone and especially with lack of arginylation resulted in endo-lysosomal compartments stress. Importantly, we provided several lines of evidence indicating that the most crucial role of Ubr1 in autophagy is to guide ubiquitinated/arginylated-cargo to SQSTM1/p62 and ensure SQSTM1/p62 oligomerization-dependent selective autophagosome/endo-phagosome maturation. These mechanisms represent a previously unrecognized Ub E3-ligase-dependent Stress QC route targeting damaged proteins for Autophagy QC. Notably, we demonstrate that damaged integral PM proteins can be substrates to Ubr1 at the ER QC and Endo-phagy QC. Ubr1 likely supports during proteostasis stress the constitutive PM-endosome PQC pathway using CHIP and ubiquitin-dependent ESCRT-machinery described here and previously^6, 7, 9^. Collectively, these findings link Ubr1 stress QC to a major biological disease pathway of Ca^2+^-signalling and selective autophagy with implications to a variety of human diseases (Summary Fig. 8H and S6F).

Multiple Ca^2+^-binding proteins link Ca^2+^-changes to ubiquitination, arginylation and biogenesis of autophagosomes and lysosomes^69^. For example, Ca^2+^-binding to the E3-ligase NEDD-4 releases its autoinhibition^70^, and CUL3 substrate recognition requires Ca^2+^-binding proteins, PEF1 and ALG2^71^. Arginylation of calreticulin and its association with cytosolic stress granules was observed only upon stressors changing cytosolic Ca^2+^ levels^72^.

Although the PM/endosomal compartment has emerged as a source of autophagic precursor membranes^22, 73–75^, the role of Ca^2+^ in autophagy is likely case-specific^76, 77^. Both Rab11+ endosomes and PI3P membranes^22^ represent a critical platform for the adaptor WIPI2, ATG16L1, ATG9 and LC3 conjugation with dynamin 2-dependent endosomal tubule scission, leading to autophagosome formation^73, 74, 78^. Rab11+ membranes were found as sites where autophagosome engulfed cytosolic mutant huntingtin exon 1 and damaged mitochondria (mitophagy) in complex with SQSTM1/p62^22^.

We expand the role of specific cargo selective endo-phagy by unravelling key steps of cargo recruitment into autophagosomes upon proteostasis stress via the conformational-sensitive arm of PQC. Here we show that both ubiquitination and arginylation in misfolded MLC1 variants by the Ubr1-chaperone complex and arginyltransferase guide SQSTM1/p62 (and LC3) to mediate endo-phagy. This selective stress autophagy mechanism is universal as a variety of arginylated Ubr1-client proteins were required to preserve the endosomal compartments health during Ca^2+^-overload. Whether Ubr1 targets are activated by Ca^2+^- and/or other stressors require investigation.

Dual post-translational modifications in the QC by organelle autophagy are not without precedent and likely serves to enhance the fidelity of structurally impaired protein elimination. Amplified mitophagy relies on both phosphorylation and ubiquitination by PINK1 and QC E3-ligase Parkin in Parkinson’s disease^14^. The alpha-1 antitrypsin variant was sequestered as a ubiquitinated and arginylated complex by SQSTM1/P62 for ER-phagy^15, 16^. Selective endo-phagy requires sequential ubiquitin-arginine modifications and is likely initiated at Rab11A+ early endosomes with the association of PI3P and Rab11A by WIPI2 and the recruitment of ATG16L1^22^. Rab11 influences cargo recycling between early and recycling endosomes, and rerouting damaged cargoes from recycling is prudent for protecting the homeostasis of the PM and late endosomes/lysosomes^58, 79, 80^.

Only a few Ub QC-ligases have been identified at the PM-endosomes or for proteostasis stress. CHIP relies on the Hsc70/Hsp90-chaperone-dependent mechanism, while the endosomal RFFL recognizes conformationally defective substrates independently of chaperones^6, 7, 9, 11^. The yeast Rsp5^12^ limits the PM accumulation of heat-stressed proteins in concert with arrestin-related adaptors^81^. Ubr1 has been invoked in chaperone-dependent and - independent^45, 65, 82^, and stress-induced cytosolic- and ER QC functions in yeast^60^. Unlike CHIP, Ubr1 has not been associated with membrane protein QC at the PM-endosomes or Stress QC in mammalians. We demonstrated that Ubr1 is a QC-ligase that has a pivotal role in the ERAD of conformationally defective MLC1s similar to a previously found role for yeast CFTR^46^ and endo-phagy QC during Ca^2+^ proteostasis stress in human cells.

The dynamics of molecular chaperones and proteotoxic stress may play a role for both CHIP and Ubr1 in QC. CHIP loss-of-function increases sensitivity to senescence, stress and ageing^83–85^. Thus, stress-activated Ubr1 can constitute an important backup QC mechanism toward certain clients. Where the Hsc70-CHIP complex recognizes substrates with exposed short hydrophobic sequences, Ubr1 as n-recognin targets destabilizing amino acids^86^. Our data and a recent report^60^ suggest alternative mechanisms for the stress PQC. Upon acute stress, CHIP can sense Hsp70 deficiency and relocate to support membrane organelles QC independently of chaperones^87^. Osmotic stress activates Ubr1 independently of the UBR-box to degrade some misfolded cytosolic and ER membrane proteins in yeast^60^ consistent with its n-recognin-independent function^65, 82^. We found that the lack of UBR-box in Ubr1 enhances QC, Ubr1 is recruited with Hsp90-complex and is a requirement for SQSTM1/p62 oligomerization, underscoring the role of Ubr1 stress QC in human cells. Diversification of Ubr1 recognition capacity during proteostasis stress may expand the limited substrate specificity and capacity of the mammalian QC ligases.

The plethora of genetic mutations, including in risk genes for cancers and neurodegenerative diseases, underscores the importance of proper protein/membrane flow at the endosomal pathway. The identified Ubr1 QC-mechanism advances our understanding of these diseases by demonstrating the capacity of cells to alleviate proteostasis Ca^2+^-stress clients toward selective auto/endo-phagy. Our results highlight the role of ubiquitination and arginylation in the stress QC and offer novel therapeutic targets in diseases afflicting membrane proteins and organelle proteostasis.

## Materials and Methods

### Experimental Models

Parental or inducible Lenti-X Tet-On ^6^ HeLa and U251N cells with and without 2HA-MLC1 (GeneID: 23209) expression were used in experiments and cultured under standard conditions^31, 32, 34^. Lentivirus production, transduction and doxycycline induction^6^ were done as described before^31^ as well as transfection of GlialCAM-Flag^31, 34^. Transfection of plasmid DNA was performed using Lipofectamine 2000 (Thermo Fisher Scientific) and siRNA using RNAiMAX or Oligofectamine transfection reagent (Thermo Fisher Scientific). Thermolabile E1 mutant CHO cells *ts20* and control E36 cells were preincubated at 40°C for 3 hr to inactivate E1^7^. Flag-Ubr1-wt in pCMV-Tag2B was a gift from Y.Yamaguchi (GeneID: 499877). C1011 was mutated to alanine to generate inactive ligase (CI). To create a mCherry vector, Ubr1-wt and Ubr1-CI were transferred to pmCherry-C1 (Clonetech).

### Immunoprecipitation and protein analyses

In total cell IP, the Ab was added directly to cell lysates. Selective isolation of MLC1-complex from the PM was achieved by cs-IP using anti-GlialCAM Ab (1:2000, R&D Systems). Ab was bound on ice to live cells for 45min, after which the unbound Ab was washed off. Cells were lysed in Triton X-100 lysis buffer (1% Triton X-100, 25 mM Tris-Cl, 150 mM NaCl, pH 8.0, 10 μM MG132 containing 20 μM PR-619, 10 μg/ml pepstatin + leupeptin, 1 mM phenylmethylsulfonyl fluoride, and 5 mM *N*-ethylmaleimide) on ice or in co-IP assays, a milder lysis buffer was used by changing Triton X-100 to 0.4% NP-40. For detecting direct ubiquitination of MLC1, lysates were denatured using 1% SDS for 5min, after which the SDS concentration was adjusted to 0.1%. The second IP step was performed using anti-HA (Biolegend, for MLC1). Treatment with cycloheximide (100 μg/ml) was carried out in full medium at 37°C for indicated times. BFA treatment (5 μg/ml) was done in the full medium at 37°C for 20h. Autophagosome-lysosome fusion was inhibited by adding Bafilomycin A1 (200 nM) and proteasomes with Bortezomib (1μM) for indicated times. Molecular chaperone Abs were anti-Hsp90, Hsc70 and Hsp40 (DNAJB1) (Enzo Lifesciences), ESCRT Abs anti-TSG101 (Santa Cruz), -Stam (Santa Cruz), -Hrs (Santa Cruz), other Abs anti-Na^+^K^+^ATPase (Abcam), -Ubr1 (Abcam), -SQSTM1 (Abcam), -LC3b (Genetex, Merck) and P4D1 anti-Ub Ab (Santa Cruz).

Cross-linking of the PM-endosomal MLC1 to Ubr1 was performed using dithiobis(succinimidyl propionate) in 0.05mM (Thermo Fisher Scientific) for 10min at room temperature and cs-IPed as above. Proximity biotinylation based on method ^88^ between MLC1 and Ubr1 was performed 24h with 50 μM biotin and cs-IP was performed as above. Eluate was diluted to Triton X-100 lysis buffer containing 0.4% SDS. Biotinylated proteins were isolated using a Streptavidin (Thermo-Fisher Scientific) pull-down assay. Beads were collected and washed with lysis buffer four times and twice with 1 ml of 2% SDS in dH_2_O. Bound proteins were removed from the magnetic beads with 50 μl of Laemmli SDS-sample buffer saturated with 20mM biotin at 98°C. For all assays, proteins were separated using SDS-PAGE and Western blot analysis. Densitometric analyses were done measuring signal intensity at the linear range using Image Studio Lite (LiCOR) or Fiji (National Institutes of Health). When quantification was done for the IP samples, the signal was normalized to the number of precipitated MLC1.

### Post-translational modification identification by mass spectrometry

Affinity purification of MLC1 was done using anti-HA in a lysis buffer as described above and final washing twice in 50 mM NH_4_HCO_3_. Beads were resuspended to 20mM Tris-HCl (pH 8.0) and 750ng of trypsin was added for 24h, and an additional 250 ng for 3h, and incubated at 37°C. Peptides were lyophilized and formic acid was added in 2%. LC-MS/MS was done on the Orbitrap Fusion Tribrid Instrument connected to an UltiMate 3000 UHPLC liquid chromatography system (Thermo-Fisher Scientific). The acquired raw files of mass spectra were searched against the Uniprot human database and analyzed using Byonic or MaxQuant with dynamic modifications for ubiquitination (+ 114.04293 Da) and arginylation (+156.10111). The false discovery rate (FDR) was set to 1%.

### Live-cell ELISA

The PM/endosomal protein expression and turnover were measured using the PM epitope labelling and cs-ELISA in live cells^6, 7, 32^. Endosomal internalization was measured for 5 min and turnover for indicated times at 37°C. TfR was measured using horseradish peroxidase (HRP)-conjugated transferrin (Thermo Fisher Scientific) and CD4 with anti-CD4 (BD Pharmigen). The transferrin-HRP or HRP-conjugated secondary Abs were measured either by luminescence using HRP-Substrate (SuperSignal West Pico, Thermo Fisher Scientific) or Ampilite (ATT Bioquest) or Amplex Red assay (Thermo Fisher Scientific).

Differences in cytosolic Ca^2+^-fold levels were measured using Calbryte 520 AM (5μM) (ATT Bioquest). The dye was loaded into cells for 45min at 37°C and 30min at RT after which 1mM Probenecid was added to each well before the fluorescence plate reading.

### Lysosomes exocytosis and enzyme assays

The lysosomal exocytosis to the PM or lysosomal beta-hexosaminidase secretion was measured in MLC1-wt or misfolded variant expressing cells. Parental and potassium channel hERG expressing cells ^6^ were used as controls. Lysosomal marker Lamp1 (Abcam) appearance to the PM indicating lysosomal exocytosis was measured by live-cell cs-ELISA. Beta-hexosaminidase secretion was monitored using a fluorometric assay. Medium or cell lysates were diluted in 0.1 M sodium citrate buffer, pH 4.5, and incubated with the 0.1mM substrate methylumbelliferyl-2-acetamido-2-deoxy-b-D-glucopyranoside (Sigma) for 1h at 37°C. The reaction was stopped by the addition of 0.5 M sodium glycine buffer, pH 10.5. Hexosaminidase activity was measured by the release of fluorescent 4-methylumbelliferone with the excitation wavelength of 360 nm and the emission wavelength of 460 nm. The total secreted enzyme was calculated from the total cellular enzyme against mg of protein. Ionomycin induction was used to increase calcium-dependent lysosomal exocytosis/secretion 5 μM for 1h at 37°C before measurements.

### Microscope imaging

The pH of endocytic vesicles containing indicated cargo molecules was measured using live-cell single vesicle fluorescence microscopy and image analysis as described previously^6, 7, 31, 34, 56, 57^. Membrane protein cargo was labelled sequentially with appropriate primary Ab against extracellular epitope and with FITC-goat anti-mouse secondary Fab (Jackson Immunoresearch). Internalization was initiated at 37°C and allowed to continue for indicated times. Recycling endosomes were labelled with 5 μg/ml FITC-Tf (Jackson Immunoresearch) for 1 h. At least >250 vesicles from 25-50 cells per experiment were analyzed, and the average weighted mean was calculated for at least three or more independent experiments. The analysis was performed on an inverted fluorescence microscope Nikon TI-E equipped with Lumencor Spectra × light source and electron-multiplying charge-coupled device (Photometrics) equipped with an Evolve 512 electron-multiplying charge-coupled device (EM CCD) camera (Photometrics Technology) and a 63×/1.4 numerical aperture (NA) Plan Apochromat oil-immersion objective. The acquisition was performed at 490 ± 5– and 440 ± 10–nm excitation wavelengths using a 535 ± 25–nm emission filter and was analyzed with NIS-Elements (Nikon).

For confocal colocalization microscopy (Leica TCS SP8X confocal microscope or LSM780 microscope, Carl Zeiss MicroImaging, 63×/1.4 NA Plan Apochromat oil-immersion objective) cells were cultured in 100 μg/ml poly-l-lysine coated coverslips. For the PM staining or endo-lysosomal cargo visualization, the PM proteins were labelled in live cells and allowed to internalize for indicated times. Cells were fixed with 4% paraformaldehyde in PBS for 15 min. Intracellular antigens were visualized in fixed, permeabilized cells using the indicated primary Abs in PBS-0.5% BSA for 1 h at room temperature. Following antibodies were used: anti-EEA1 (Cell Signalling), -Flag (M2, Sigma), -ERp57 (GeneTex), -LAMP1 (Abcam, Novus Biologicals), -LAMP2 (Abcam), -CLC (Sigma), -AP2 (Sigma), Ubr1 (Abcam), -SQSTM1 (Abcam), -LC3b (Genetex, Merck), -R-Bip (Merck), -Arg (Thermo Fisher Scientific), Phalloidin-Alexa 594 (Thermo Fisher Scientific). Secondary antibodies were from Thermo Fisher Scientific (Anti-Mouse Alexa 488, Anti-Rabbit Alexa 555, Anti-Rabbit Alexa 488, Anti-Goat Alexa 647) or Jackson Immunoresearch (Anti-Rat Alexa 647, Anti-Rat Dylight 405)

Proximity ligation assay was performed using Duolink-technology following the manufactures instructions. For endosomal detection, anti-HA Ab (Biolegends, for MLC1) was allowed to internalize in live cells. For detecting cell surface MLC1, cells were fixed before Ab binding. After MLC1 internalization or cell surface labelling, intracellular proteins Ubr1 and EEA1 were detected from fixed samples. Anti-mouse plus and anti-rabbit minus probes were used to crosslink the desired two epitopes and visualized them as described above. Manders’ overlap coefficient was quantified from 25-200 cells and Lamp+ lysosomal diameter from >2000 organelles using Fiji-plugins.

Cells on glass coverslips were loaded with Fura2-AM (5 μM, Thermo Fisher Scientific) for 30min, washed and incubated in Krebs–Ringer–Hepes solution for 10min at 37 °C. The signal was measured using a Nikon TE300 Eclipse microscope equipped with a Sutter DG-4/OF wavelength switcher, Omega XF04 filter set for Fura-2, Photonic Science ISIS-3 intensified CCD camera and MetaFluor software. Images were obtained every 20s using a 20X objective. Ratio values (340/380 nm) were transformed to cytosolic [Ca^2+^] using the equation derived by^89^.

### Statistics

Paired or unpaired two-tailed Student’s *t*-test was used for p-values as indicated in the figure legends. Statistical significance was set to p< 0.05. All data in curves and bar plots represent the means average of at least three or more independent experiments. Data are expressed as means ± SEM.

## Acknowledgements

We thank Yong Tae Kwon (Seoul National University, Republic of Korea), Cheol-Sang Hwang (POSTECH, Republic of Korea) for materials and technical advice; Masayuki Komada, Yuki Yamaguchi (Tokyo Institute of Technology, Japan) and Alexander Varshavsky (California Institute of Technology, USA) for technical advice; Andrew Marshall (Coherent Scientific, Australia) with Nikon Ti and NIS elements; Monash Biomedical Proteomics Facility for providing instrumentation and technical support.

## Abbreviations

α: anti
CHIP: C-terminus of Hsc70-Interacting Protein
cs: cell surface
ER: endoplasmic reticulum
ERAD: endoplasmic reticulum-associated degradation
ESCRT: endosomal sorting complex required for transport
GlialCAM: glial cell adhesion molecule
IP: immunoprecipitation
LAMP1: lysosomal associated membrane protein 1
MLC1: megalencephalic leukoencephalopathy with subcortical cyst
ns: non-significant
NT: non-target
PM: plasma membrane
PQC: protein quality control
siRNA: small interfering RNA
Ub: ubiquitin
Ubr1: ubiquitin-protein ligase E3 component n-recognin 1.

## Disclosure Statement

The authors declare no competing interests.

## Additional information

### Author Contributions

B.B.W, S.I, H.X and G.Y.R conducted experiments and P.M.A. compiled figures. C.H and R.B.S performed LC-MS/MS. X. E.-V and R.E provided materials. G.L.L provided materials, scientific advice and gave critical input regarding the manuscript. P.M.A wrote the manuscript.

## Notes

### Competing Interest Statement

The authors have declared no competing interest.

