## Supplemental material for "Ubr1-induced selective endo-phagy/autophagy protects against the endosomal and Ca^2+^-induced proteostasis disease stress"

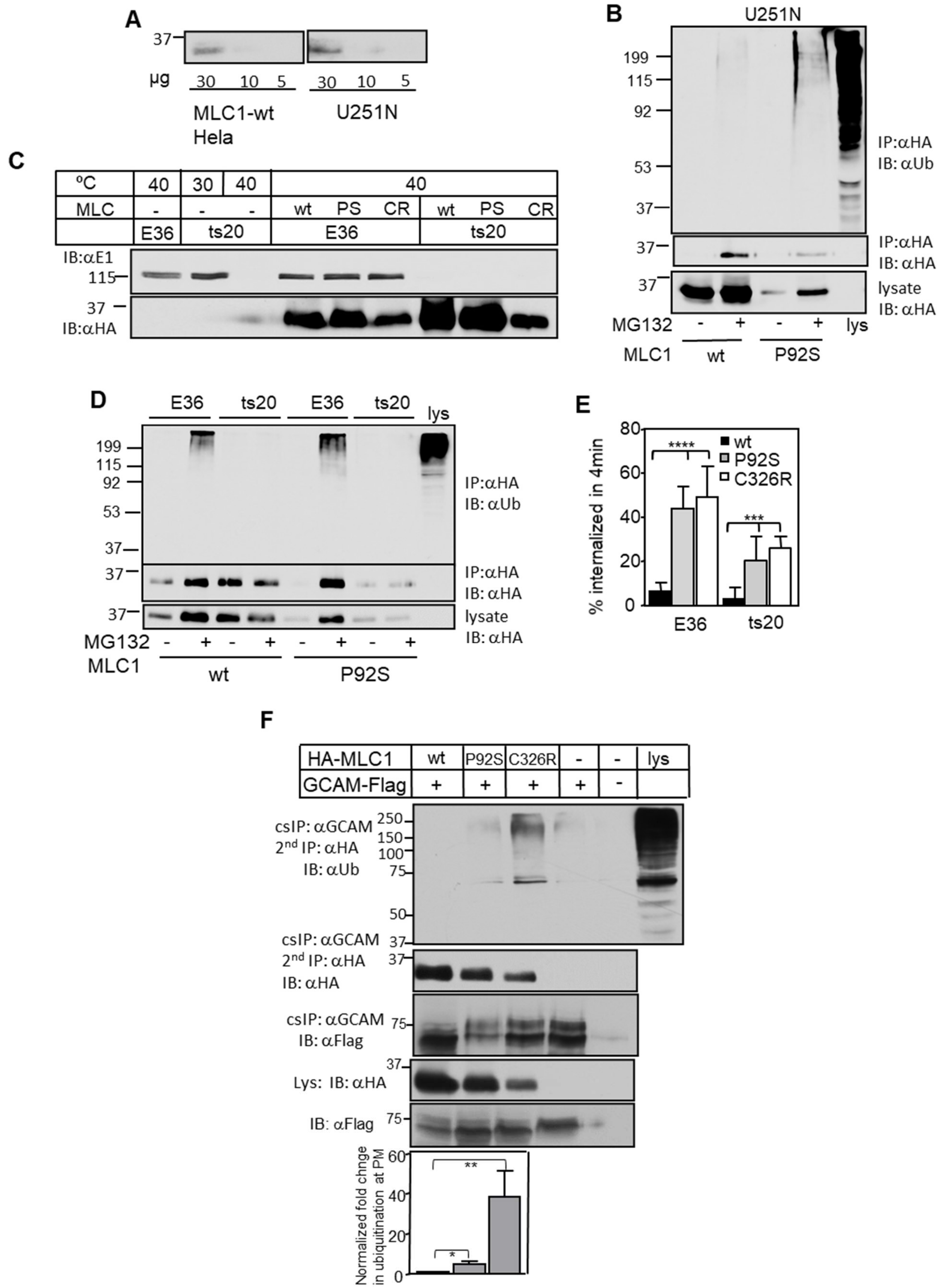

A
MLC1-wt Hela
U251N
B
U251N
IP:αHA IB: αUb
IP:αHA IB: αHA
lysate IB: αHA
MG132
MLC1 wt P92S
lys
C
ºC
MLC
wt PS CR
E36 ts20
IB:αE1
IB:αHA
D
E36 ts20 E36 ts20 lys
IP:αHA IB: αUb
IP:αHA IB: αHA
lysate IB: αHA
MG132
MLC1 wt P92S
E
% internalized in 4min
wt
P92S
C326R
E36 ts20
F
HA-MLC1 wt P92S C326R lys
GCAM-Flag
csIP: αGCAM 2nd IP: αHA IB: αUb
csIP: αGCAM 2nd IP: αHA IB: αHA
csIP: αGCAM IB: αFlag
Lys: IB: αHA
IB: αFlag
Normalized fold chnge in ubiquitination at PM

**Figure S1. Astrocyte membrane regulatory cluster MLC1 disease variants are ubiquitinated at the PM-endosomes**

**A)** Immunoblotting of HA-MLC1-wt expression in stable inducible HeLa to endogenous MLC1 in astrocytic U251N cells using an anti-MLC1 antibody. The lysate amount is indicated. **B)** Misfolded MLC1 is more ubiquitinated than the wt in astrocytic U251N cells. Immunoprecipitation with anti-HA was performed for MLC1-wt and P92S mutant in denaturing conditions to detect direct ubiquitination of MLC1. Degradation of MLC1 was inhibited with MG132 for 2h to increase the detection of ubiquitination. Stable HA-MLC1 astrocytic U251N cells were used. Lys, lysate. **C-D)** The depletion of the temperature-sensitive (ts) E1 ubiquitin-activating enzyme impeded the MLC1 ubiquitination and decreased PM turnover. HA-MLC1-wt and disease associated mutations P92S (PS) and C326R (CR) were transiently expressed in E36 control and ts20 cells and the E1 was inactivated at 40ºC for 3h, and the total protein expression was analyzed by Western blotting (C). Ubiquitin conjugates in MLC1 and inhibition of degradation with MG132 were done as in A (D). Cs-ELISA was performed to detect changes in the endosomal internalized (4min) amount of MLC1 (E). **F)** The PM-endosomal MLC1 misfolding increases ubiquitination. The MLC1/GlialCAM complex was first cell surface immunoprecipitated (cs-IP) using GlialCAM Ab and the second IP was performed with HA Ab for MLC1 after denaturation. GlialCAM-Flag was transiently expressed in stable inducible HA-MLC1 HeLa. The ubiquitination was normalized to the MLC1 amount at the PM. GCAM, GlialCAM; lys, lysate; α, anti. Means ± SEM, n≥3. p-value: *<0.05, **<0.01, ***<0.001 ****<0.0001.

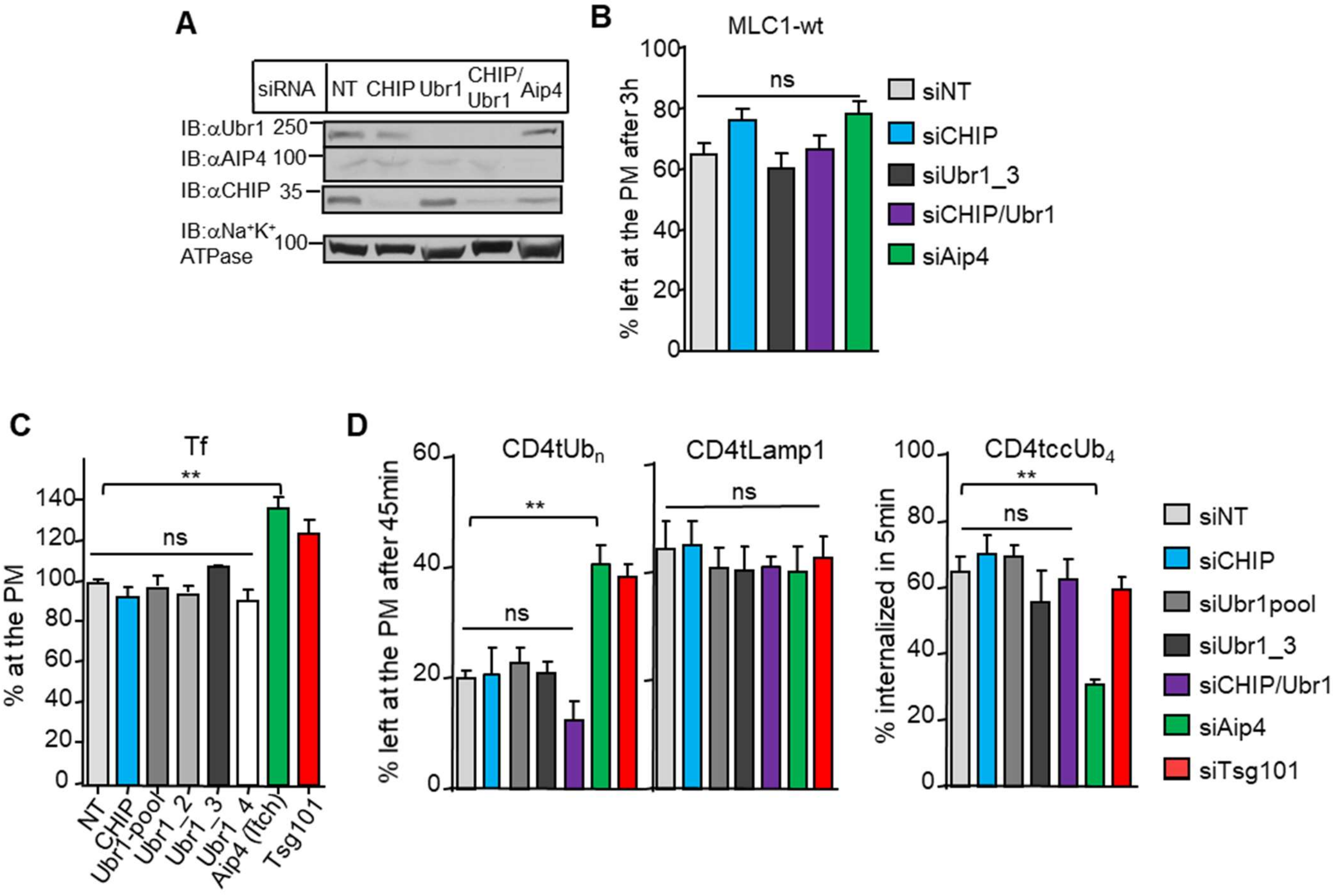


**Figure S2. CHIP and Ubr1 are post-Golgi QC E3 ubiquitin ligases**

**A**) Western blot analysis of siRNA cells from Fig.2E depleted for candidate QC E3 ligases. (**C**) Candidate E3 ligase depletion effect on native HA-MLC1-wt stability at the PM was measured using cs-ELISA expressed in stable inducible HeLa. Differences are not statistically significant (ns). (**D**) Candidate E3 ligase depletion effect on cell recycling endosomes. Internalization was measured for transferrin receptor (Tf) using cs-ELISA. SiNT served as a negative and siTsg101 as a positive control. Only Aip4 (Itch) had a significant effect. (**E**) Candidate E3 ligase depletion (siRNA) effect on constitutively polyubiquitinated ($CD4tUb_n$), lysosomal targeted (CD4tLamp1) or tetra-ubiquitinated cargo ($CD4tccUb_4$) in stable HeLa. The PM turnover was measured for $CD4tUb_n$ and CD4tLamp1, and endosomal internalization (5min) for $CD4tccUb_4$ due to its fast turnover. Only Aip4 (Itch) had a significant effect on ubiquitinated cargo ($CD4tUb_n$ and $CD4tccUb_4$) when compared to NT control indicating non-specific ubiquitinated cargo recognition. NT, non-target, Means ± SEM, n≥3. p-value: *<0.05, **<0.01, ***<0.001 ****<0.0001.

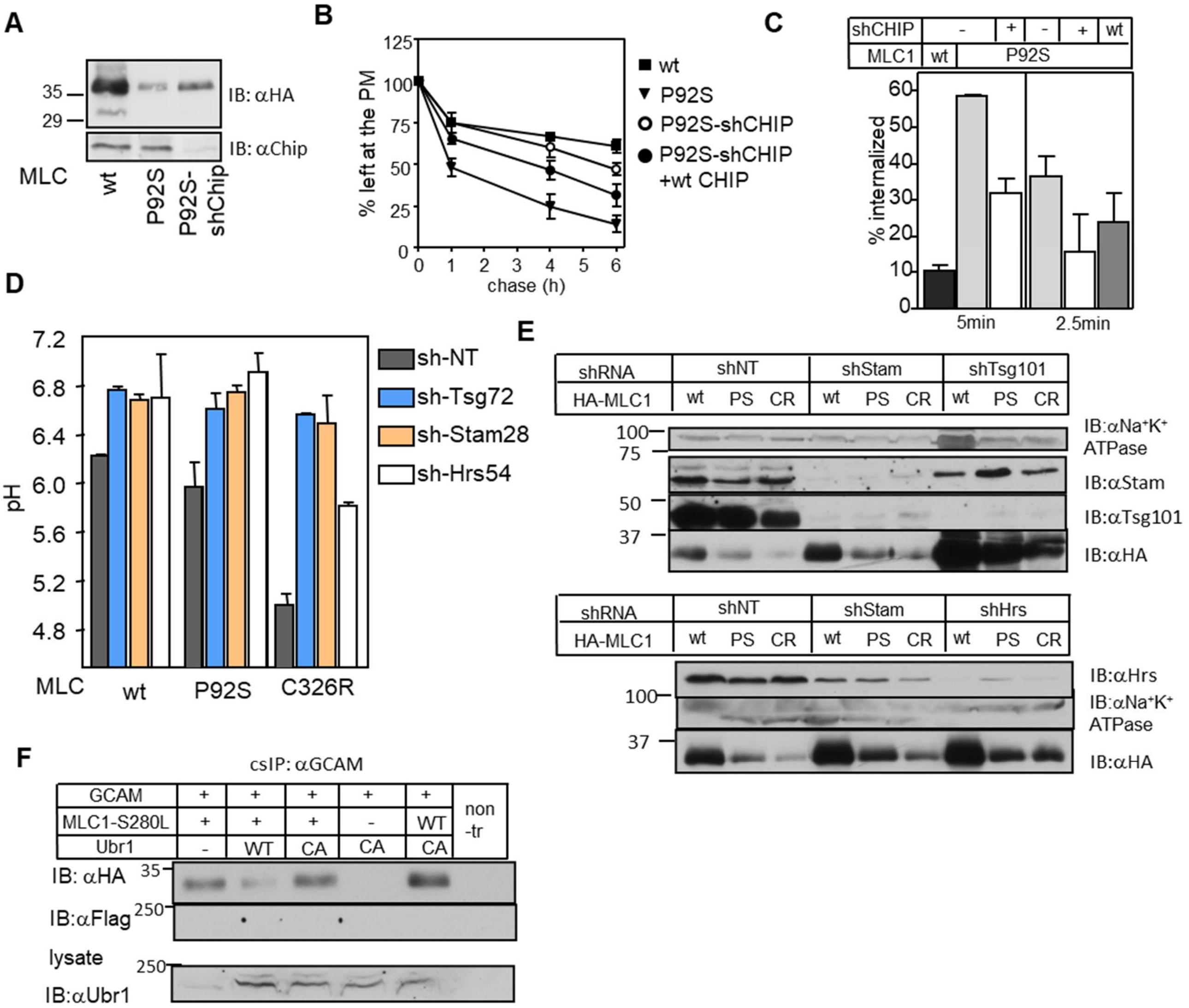


**Figure S3. Classical CHIP and ESCRT-mediated ubiquitin-dependent endo-lysosomal QC pathway**

**A-C**) The depletion of molecular chaperone-dependent E3 ubiquitin ligase CHIP decreased the PM turnover of misfolded MLC1. shNT or shCHIP cells were transiently transfected with disease associated with mutant P92S or MLC1-wt. Western blot analysis of total cellular lysates was done to detect total protein expression (A), and cs-ELISA to detect changes in the PM turnover (B) and internalization (C). **D**) The depletion of ESCRT0-I components impaired the lysosomal targeting of misfolded MLC1 mutants. Vesicular pH measurement was performed for MLC-wt or disease associated mutant P92s or C326R containing endosomes. Vesicular pH of MLC1 cargo containing endosomes was determined in cells depleted for Tsg101, Stam2 or Hrs or control shNT. Mean vesicular pH after 1h chase is shown. The values are means ± SEM of 3 independent experiments. **E**) Western blot analysis of shESCRTs and shNT cell lines used in I for MLC1-wt or misfolded mutants P92S or C326R. (**F**) Association of Ubr1 with misfolded MLC1-S280L. Cells expressing S280L mutant were overexpressed with Flag-Ubr1-wt or CA. Cross-linking was done with dithiobis(succinimidyl propionate), after which the cs-immunoprecipitation of the MLC1/GlialCAM -complex was performed. Ubr1 association with S280L was seen faintly in lanes 2-3. MLC1-wt was used as a folded native control.

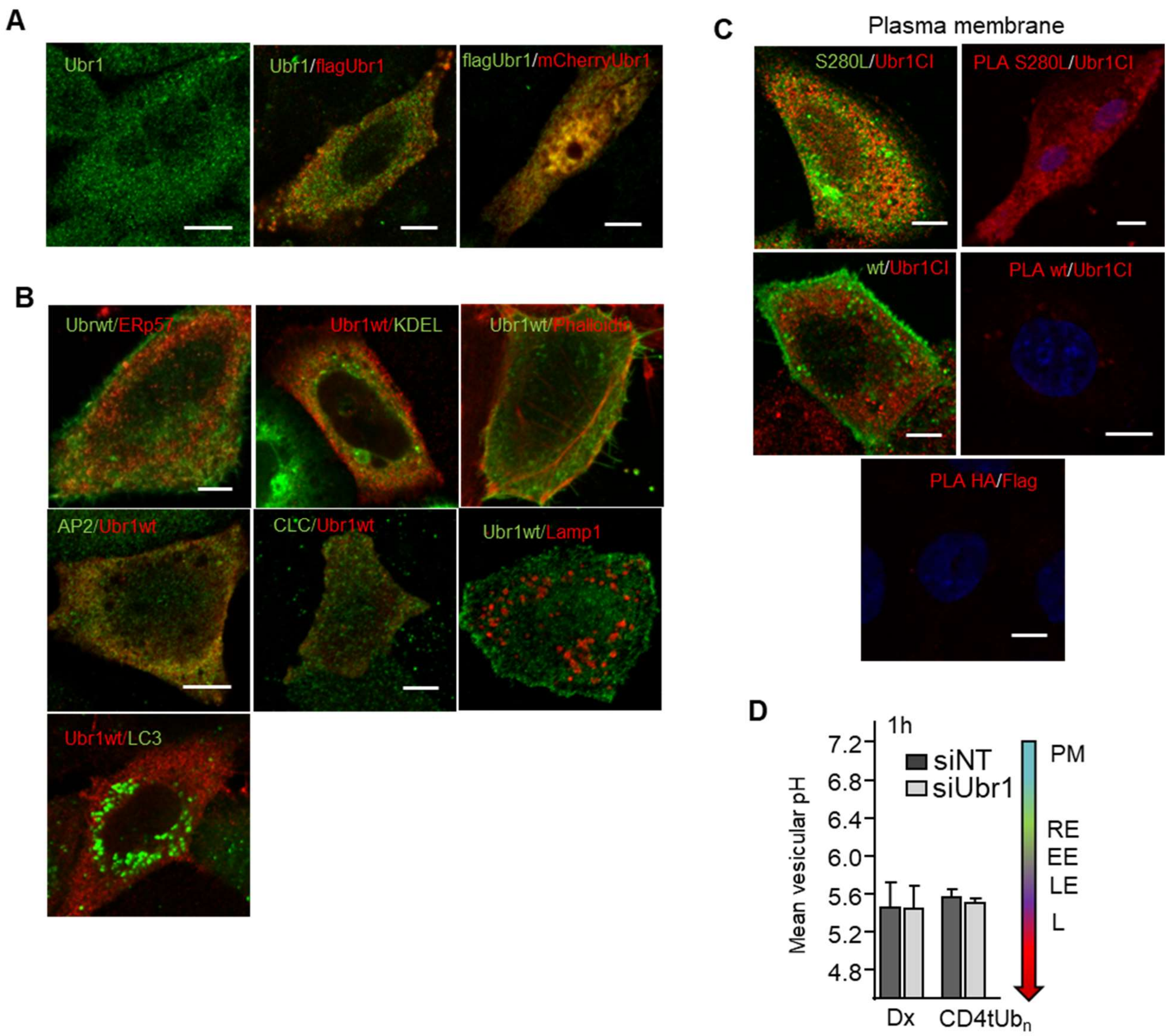


**Figure S4. Subcellular location of Ubr1**

**A**) Expression of Ubr1 was detected by using anti-Ubr1 (endogenous) or expressing Flag-Ubr1-wt or mCherry-Ubr1-wt. The appearance of nuclear Ubr1 varied and may be cell cycle-dependent. Bar 5 µm. **B**) Ubr1 expression in different subcellular markers was monitored using Flag-Ubr1-wt and markers for the ER (anti-ERp57 or GFP-KDEL), F-actin at endosomal invagination sites (phalloidin), endosomal clathrin-coated vesicles (AP2 and CLC), lysosomes (Lamp1) or autophagy (GFP-LC3). Bar 5 µm. **C**) Interaction of MLC1-S280L and Ubr1 at the PM. The S280L at the PM was labelled without internalization with flag-Ubr1-CA, and PLA was performed between HA-Flag epitopes. MLC1-wt and cells without S280L expression were used as a control. Bar 5 µm. **D**) Controls for Ubr1 specificity toward misfolded MLC1 (as in Fig.S2). Fluid phase marker dextran (Dx) and poly-ubiquitinated model ($CD4tUb_n$) were used as control cargoes for siUbr1. Mean vesicular pH measurement of cargo labelled endosomes was determined using live-cell microscopy in cells depleted for Ubr1 or control siNT. PM; plasma membrane, RE; recycling endosome (pH 6.3-6.2), EE; early endosomes, LE; late endosome, L; lysosome.

A

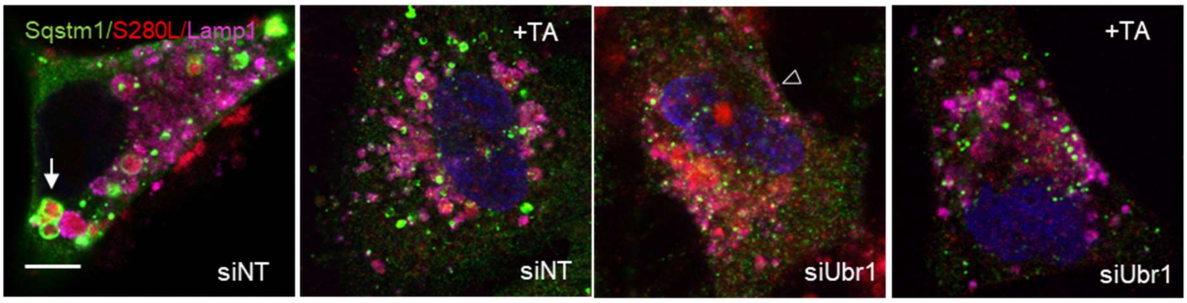


B

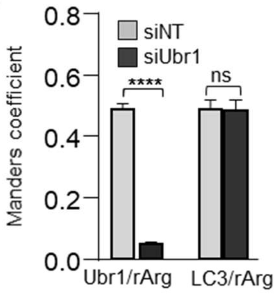


C

| MLC1 | identified peptide | PTM |
|---|---|---|
| N-term | R.GRQDPASYAPD+[+156.1011]AK[+114.04]PSDLQLSK.R | Ub, R |
| | R.QDPASYAPD[+156.1011]AK[+114.04]PSDLQLSKR[+156.1011].L | Ub, R |
| 1st loop | TV.SRRNANVIPN FQ | none |
| 2nd loop | R.SSE[+156.1011]E[+156.1011]DCKK[+114.043].K | R, R, Ub |
| | K.K[+114.043]GSMSDSANILDEVPFPAR.V | Ub |
| | K.GSMSDSANILDEVPFPAR | none |
| | K.GSMSDSANI[+156.1011]LD[+156.1011]EVPFPAR.V | R, R |
| 3rd loop | AA ECPSKCLVEV | none |
| C-term | R.FK[+114.043]VSAR.L | Ub |
| | R.SPLK[+114.043]EFDK.E | Ub |
| | .E[+156.1011]FD[+156.1011]KEKAWR.A | R, R, |

**Figure S5. Proteotoxic stress induces Ubr1, ubiquitination, arginylation and autophagy to safeguard endo-lysosomal pathway proteostasis**

**A**) The connection of Ubr1 to arginylation and SQSTM1/p62 recruitment in MLC1-S280L expressing HeLa cells. When indicated, tannic acid (TA) was used to inhibit arginylation and Lamp1 to mark lysosomes. The arrow indicates autolysosomes/autophagosomes and arrowheads lysosomal Lamp1 at the PM. Bar 5 µm. **B)** Manders' overlap coefficient of Ubr1 and arginylated substrates (rArg) (representative figure 7A) and autophagy marker LC3b with arginylated substrates (rArg) (representative figure 7B) in MLC1-S280L U251n cells. n>47 cells, ****p<0.0001, ns; non-significant. **C**) Mass spectrometry analysis of affinity-purified MLC1-wt and S280L amino acids at the intracellular part for arginylation (R) and ubiquitination (Ub). Intracellular peptide fragments, their location in the MLC1 and modifications are indicated. The schematic figure is in 7C.

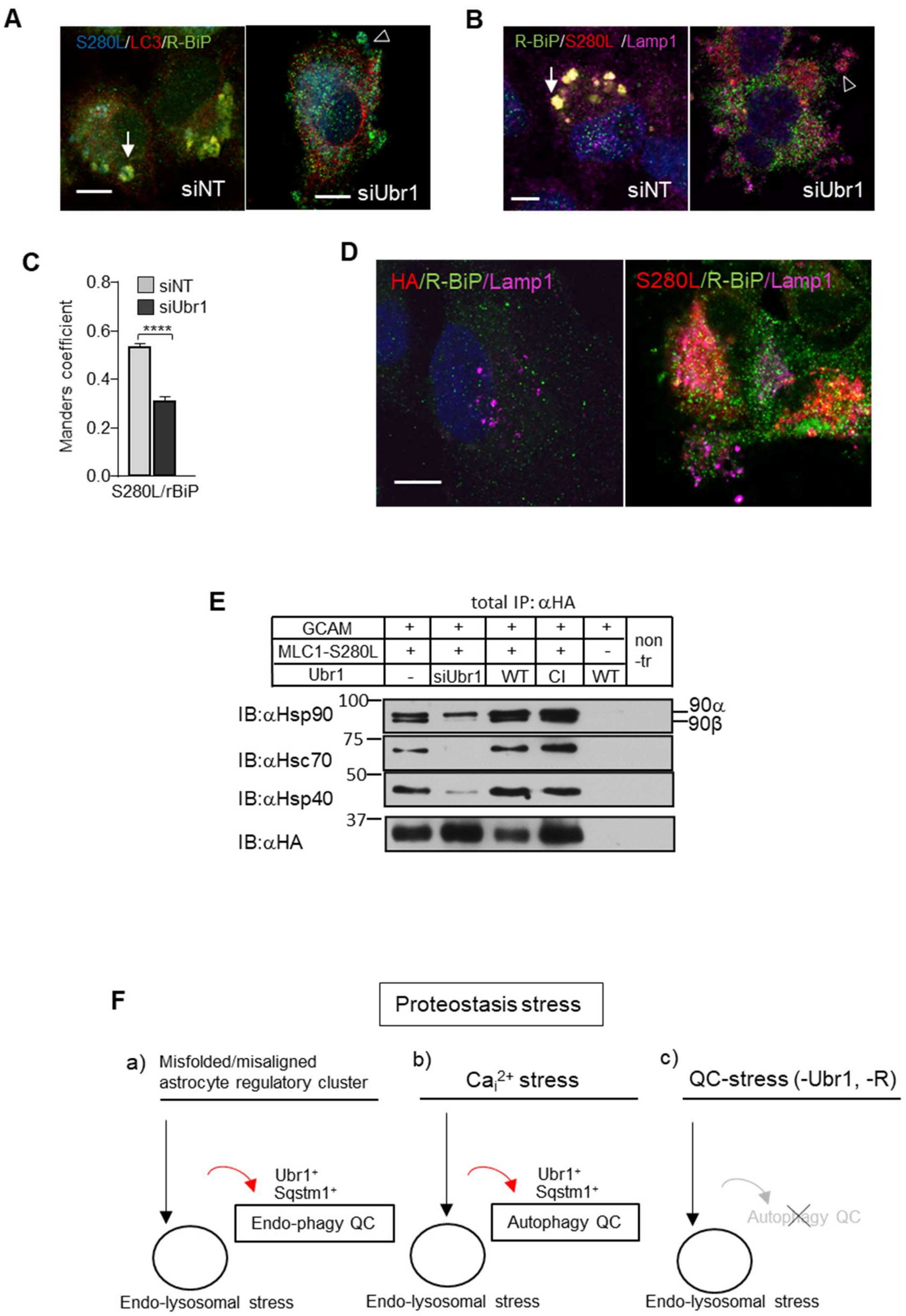


**Figure S6. Ubr1 is a proteotoxic stress-induced Endo-phagy and Autophagy QC ligase**

**A**) Autophagy marker LC3b and R-BiP colocalization analysis in MLC1-S280L expressing U251N cells depleted for Ubr1 or siNT. The PM labelled MLC1 was internalized for 30min to identify the endosomal S280L location. Arrow points autolysosomes and arrowhead points the PM blebs with LC3b. Bar 5 µm. **B)** Lysosomal marker Lamp1 and R-BiP colocalization analysis in MLC1-S280L expressing U251N cells depleted for Ubr1 or siNT (as in A). Arrow points autolysosomes and arrowheads the PM blebs with Lamp1. Bar 5 µm. **C)** Manders' overlap coefficient of R-BiP and S280L colocalization in U251N cells treated with siNT or siUbr1. Representative figures are in A-B. n>37 cells, ****p<0.0001. **D)** MLC1-S280L increases arginylated BiP retrotranslocation to the endo-lysosomes. Lysosomal marker Lamp1 and arginylated BiP was analyzed in MLC1-S280L and control U251N cells. Bar 5 µm. **E)** Association of molecular chaperones in total cell lysates with MLC1-S280L in Ubr1 (related to figure 7E). Cells were depleted for Ubr1 or overexpressed with flag-Ubr1-wt or CA. Coimmunoprecipitation was done using anti-HA and probed against indicated molecular chaperones. **F)** Summary of Ubr1 functions during cellular proteotoxicity to fine-tune membrane organelle and PQC in $Ca^{2+}$-stress and disease by using endo-phagy and autophagy pathways. Ubr1 regulates the rerouting of misfolded/misaligned astrocyte regulatory protein MLC1-cluster causing endosomal compartment proteotoxicity to Endo-phagy QC (a), and substrates during $Ca^{2+}$-induced proteotoxicity to Autophagy/Endo-phagy QC (b) by alternative ubiquitin/arginine-mediated and autophagosome scaffold protein SQSTM1/P62 directed pathway. Lack of functional Ubr1 and/or argininetransferase activity result in QC-stress (c) suggesting that the Ubr1/ SQSTM1/P62 pathway is an important safeguard mechanism also in physiological conditions.
